# Excitatory delay-coupling explains in-phase and antiphase functional connectivity

**DOI:** 10.64898/2026.07.17.739148

**Authors:** Ahmet Omurtag, Andrei Dragomir, Jonathan Crofts, William W Lytton

## Abstract

Coordinated oscillatory activity between brain regions underpins cognition, yet the phase relationships governing this coordination remain poorly understood. Using scalp EEG from 31 participants performing a motor learning task, we show that inter-site phase clustering occurs predominantly at in-phase or antiphase relationships, with the transition between them governed by conduction delay. Homologous interhemispheric pairs remain in-phase despite long distances, consistent with faster callosal conduction. A minimal model of two delay-coupled excitatory populations reproduces these features without parameter tuning; stability analysis shows that in-phase and antiphase oscillations arise from competing instabilities, with delay determining which dominates. Task performance shifts a frontoparietal network toward in-phase connectivity, a modulation captured by the Phase Relationship Index (PRI) but missed by conventional clustering metrics. Preliminary evidence from independent datasets suggests these patterns generalise. These findings challenge the assumption that zero-lag connectivity reflects volume conduction, offer a mechanistic account linking conduction delay to phase organisation, and introduce PRI as a metric sensitive to functionally relevant connectivity changes invisible to existing measures.

## Introduction

Coordinated oscillatory activity across neural networks is a pivotal level of organization, connecting phenomena ranging from single neuron activity to behaviour. Its study provides an interdisciplinary framework spanning psychophysics, cognitive psychology, neuroscience, biophysics, computational modelling and mathematics^1^.

Research has revealed that inter-area phase synchronisation, including same-frequency phase synchronisation and phase-amplitude coupling, play fundamental roles across a wide range of cognitive functions: perceptual binding and feature integration^2–4^, selective attention ^5,6^, working memory maintenance ^7,8^, long-term memory encoding and retrieval^9,10^, and spatial navigation^11,12^.

A parallel body of work approaches neural synchronisation from a computational modelling and dynamical systems perspective. Studies using reduced phase models have shown how network topology, coupling strength, and transmission delays determine synchronisation regimes^13–15^. Extending this to biophysically detailed models, bifurcation analyses have further revealed transitions between fixed points, limit cycles and chaos^16,17^.

Inter-area phase synchronisation involves the alignment of oscillatory phases at the same frequency between two brain regions such that an approximately constant phase difference is maintained. Phase synchrony between regions is thought to operate independently of local firing rate, offering a distinct channel for neural information processing that takes advantage of rapid fluctuations in responsiveness^18–21^. It is often quantified by metrics known as Phase Locking Value (PLV)^22^ or Inter-Site Phase Clustering (ISPC)^23^.

Given its growing use in basic research and as a biomarker for clinical conditions, it is important to understand what phase synchronisation metrics capture and what they may discard. Zero-lag synchronisation between distant regions has sometimes been viewed with scepticism due to unavoidable axonal conduction delays, with reviewers characterising the prevailing view as a “conundrum”^4^, “problem”^24^ or “paradox”^25^. This has motivated the development of metrics designed specifically to minimize the contribution of zero-lag connections^26–29^. Yet mounting evidence indicates that zero-lag synchronization between distant regions is genuine and potentially a powerful biomarker^30,31^. Furthermore, in the dynamical systems literature, zero-lag and π -lag synchronisation are well-established as dominant forms of interaction for delay-coupled oscillators^17,32,33^.

In the present study we use current-density transformed scalp EEG from participants performing a motor learning task to show that when inter-site phase clustering is sufficiently strong, it occurs predominantly at in-phase or antiphase relationships. The transition from in-phase to antiphase is governed by cortical distance, but more precisely by conduction delay, as homologous interhemispheric pairs, despite long distances, remain in-phase presumably due to faster callosal conduction. A minimal model of two delay-coupled excitatory populations reproduces these features, and linear stability analysis reveals that the transition arises from two competing instabilities exchanging dominance as delay increases. Despite its simplicity, the model captures multiple features of the empirical data without parameter tuning and makes additional testable predictions. We also show that task performance selectively modulates a frontoparietal network to shift toward in-phase connectivity, and this modulation is captured by the Phase Relationship Index (PRI), a metric we introduce here, where ISPC alone shows little change. Preliminary results from independent datasets suggest these findings generalise beyond the present sample.

## Results

We analysed resting-state and task-related EEG from 31 participants performing a laparoscopic motor learning task (see Methods). EEG was recorded from 19 scalp electrodes and transformed using the surface Laplacian to attenuate volume conduction. For each of the 171 electrode pairs, we computed inter-site phase clustering (ISPC) and the Phase Relationship Index (PRI) within narrow frequency bands (1 Hz width). ISPC quantifies the strength of phase clustering, ranging from 0 (no consistent phase relationship) to 1 (perfect phase locking). PRI quantifies the nature of the phase relationship, ranging from 0 (in-phase) to 1 (antiphase). The experimental session comprised alternating rest and task episodes (R1, T1, R2, T2, R3, T1R, R4), with rest episodes lasting 2 minutes each.

### Phase relationships cluster toward in-phase and antiphase

Examining representative electrode pairs from a single participant centred at 16 Hz (1 Hz wide band) revealed three qualitatively distinct patterns: strongly clustered near zero phase lag, strongly clustered near π phase lag, and weakly clustered with no consistent phase relationship (**Fig. 1a-c**). An inset at the top of each column (a-c) in the figure shows a representative time segment from the pair of band filtered EEGs. For a clearer visual illustration of example phase relationships, longer time segments are provided in **Fig. S2b**. Furthermore, in addition to the three pairs shown here, a large subset of electrode pairs is provided in **Fig. S1**. The frequency band at 16 Hz was chosen for illustration; as shown later (**Fig. 2**), the results are robust across frequencies.

**Fig. 1.**
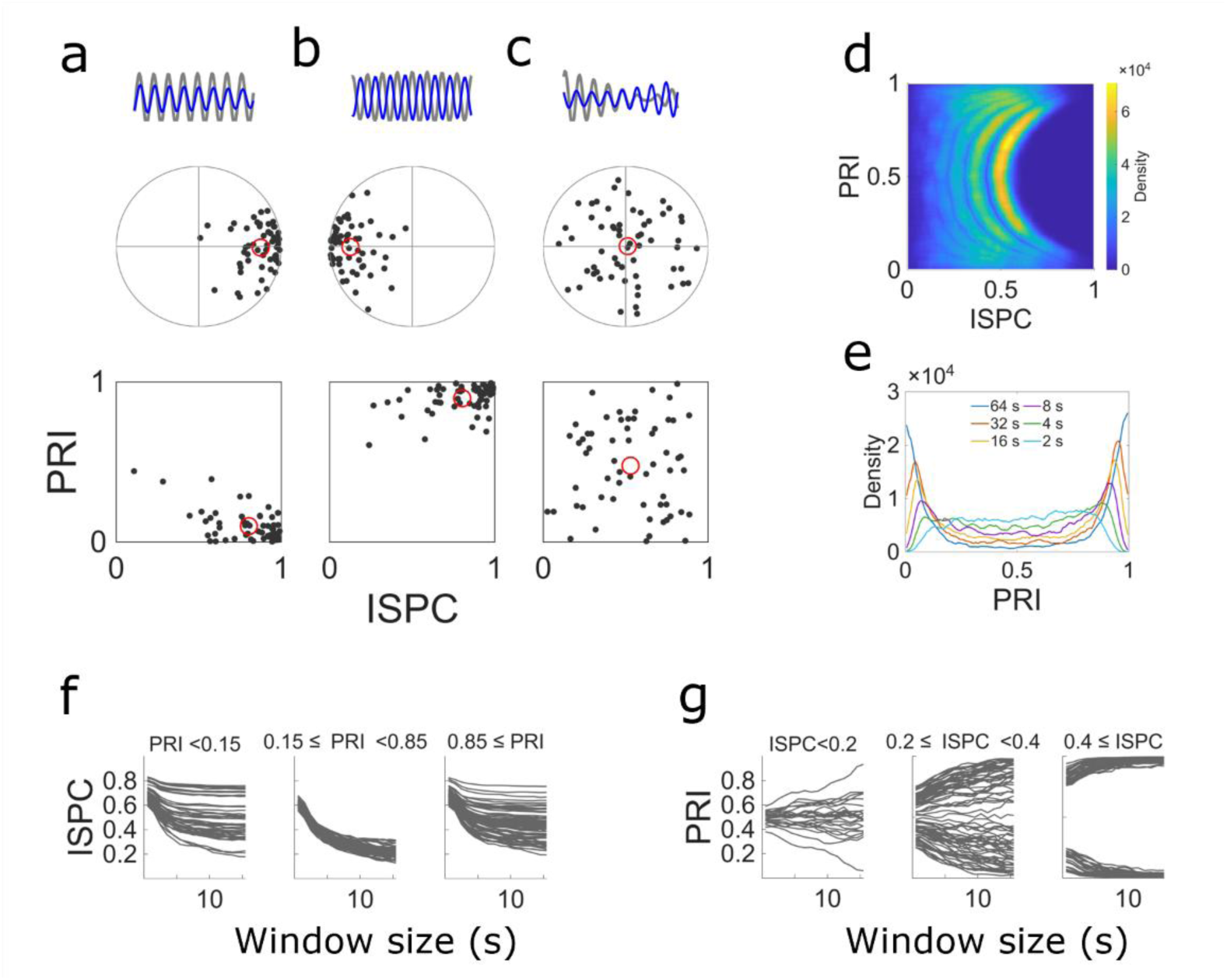
Phase clustering and its dependence on phase relationship and time window. **a–c,** Phase clustering in three representative electrode pairs from one representative participant during resting-state EEG (2 min), narrow-band filtered at 16 Hz (1 Hz bandwidth). For each pair, the inter-site phase clustering statistic *z* was computed in adjacent windows of size *W* = 2 s, yielding 60 values (black dots) with their mean (red circle). Results are shown in the Argand plane (unit circle in grey) and the ISPC–PRI plane. An inset at the top of each column shows a representative time segment from the pair of narrow band filtered signals (blue and grey). The examples show (a) the electrode pair P3–O1 that was strongly in-phase clustered; (b) Fz–T6, strongly antiphase clustered; and (c) Pz–O2, weakly clustered or unclustered. **d**, Heatmap showing the joint distribution of ISPC and PRI across all electrode pairs (*n* = 171) and participants (*n* = 31), computed separately for each window size. Each arc-shaped distribution corresponds to a different *W* (2 s at the right; then, toward left 4, 8, 16, 32, and 64 s). **e**, Marginal distribution of PRI across all pairs and participants at each window size, showing the progressive separation of in-phase and antiphase modes as *W* increases. **f**, Dependence of ISPC on *W* for low PRI (left panel), moderate PRI (middle), and high PRI (right) electrode pairs. **g**, Dependence of PRI on *W* for low ISPC (left panel), moderate ISPC (middle), and high ISPC (right) electrode pairs.

**Fig. 2.**
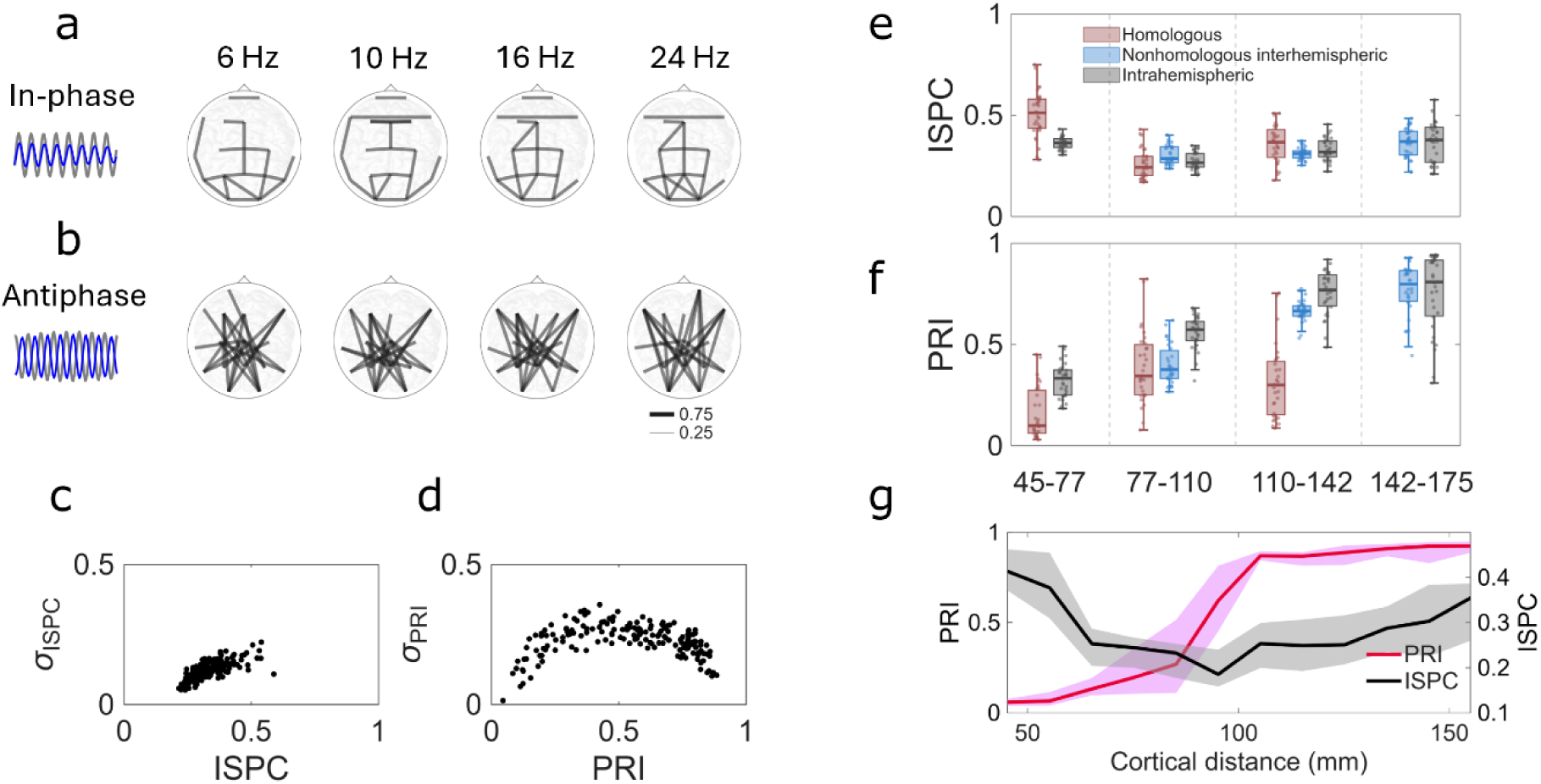
Spatial organisation, reliability and distance-dependence of in-phase and antiphase connectivity. **a–b,** Topographic maps of connections at four frequencies (labelled at top), showing electrode pairs whose participant-averaged PRI ranked in (**a**) the bottom 12% (in-phase clustering) or (**b**) the top 12% (antiphase clustering). The connector line width and transparency depend on ISPC value (see legend lower right). Inset left: a representative 0.5 s segment of narrow-band filtered EEG (16 Hz) from a typical pair, with signals from the two sites shown in blue and grey. **c,** Standard deviation across participants of the frequency averaged ISPC, with each point representing one electrode pair. **d,** As in c, but for PRI. **e–f,** Distribution of connectivity measures as a function of cortical distance, shown separately for homologous interhemispheric (red), non-homologous interhemispheric (blue), and intrahemispheric (grey) pairs. (**e**) ISPC. (**f**) PRI. **g,** Distance-dependence of phase clustering at 16 Hz. ISPC (black) and PRI (red) are plotted against cortical distance (10 mm bins). Solid curves show participant medians and shaded regions indicate the 25^th^-75^th^ percentile range across subject medians.

Across all electrode pairs and participants, the joint distribution of ISPC and PRI formed a characteristic arc-shaped pattern. Pairs with high ISPC concentrated near PRI values of 0 or 1, while pairs with low ISPC showed scattered intermediate phase relationships (**Fig. 1d**). This bimodal structure was not assumed in advance but emerged from the data. As the analysis window size W increased, the arc extended leftward, with pairs having weak clustering tendency drifting to lower ISPC, while the tips of the arc (strongly clustered in-phase and antiphase pairs) remained stable (**Fig. 1d**). The marginal distribution of PRI showed progressively sharper separation of in-phase and antiphase modes with increasing *W* (**Fig. 1e**).

To examine how ISPC and PRI stabilise with increasing window size, we plotted each measure as a function of *W* for a representative participant, grouping electrode pairs by their PRI or ISPC values. Pairs with low or high PRI, corresponding to in-phase or antiphase clustering, maintained high ISPC as *W* increased, while pairs with intermediate PRI showed ISPC that declined rapidly, reflecting their lack of a consistent phase relationship (**Fig. 1f**). Conversely, pairs with high ISPC converged quickly to either low or high PRI values, indicating stable in-phase or antiphase relationships, whereas pairs with low ISPC showed PRI values that remained scattered or converged slowly (**Fig. 1g**). The groupings of pairs into low, medium or high ISPC (or PRI) in these subplots were based on the largest window size shown, but using other windows (with suitably adjusted cutoff values) did not change the results as the relative ordering of pairs remained approximately constant with *W*. Together, these results confirm that ISPC and PRI jointly characterise a stable property of electrode pairs: strongly clustered pairs lock into either in-phase or antiphase relationships, while weakly clustered pairs exhibit no preferred phase difference.

We statistically tested the observation that phase relationships congregate near in-phase and antiphase (**Fig. 1a-c**, also **Fig. S1**). Using a doubling transformation that maps both 0 and *π* to the same point on the unit circle, we applied the V-test for circular uniformity to the phase angle *θ* = Arg(*z*), pooled across subjects, for each frequency-electrode pair combination. For window sizes *W* = 4, 8, 16, and 32 seconds, between 99.6% and 100% of combinations showed significant clustering toward 0 or π across all episodes, with FDR correction making no practical difference, indicating overwhelmingly small p-values. Task episodes (T1, T2, T1R) showed a small but consistent increase in the fraction of significant combinations relative to resting episodes (R1, R2, R3, R4). For *W* = 2 seconds, the fraction of significant combinations dropped to 53%, indicating that this window length is too short for the underlying phase relationship to establish itself reliably. This provided an empirical lower bound on the timescale of clustering, lying between 2 and 4 seconds. These results provided strong statistical support for the assertion that inter-site phase differences are directed toward either 0 or *π* whenever the phase clustering vector is estimated over a sufficiently long window.

### Spatial organisation of phase relationships

We next examined the spatial organisation of in-phase and antiphase connections. Topographic mapping revealed that electrode pairs with predominantly in-phase clustering (lowest 12% of PRI values, selected to provide a large enough sample without visual clutter) and those with predominantly antiphase clustering (highest 12% of PRI values) had distinct spatial distributions, with in-phase connections concentrated among nearby pairs and antiphase connections more prevalent among distant pairs (**Fig. 2a-b**). This pattern was consistent across all examined frequencies from 2 to 32 Hz (the figure shows four representative frequencies).

Both ISPC and PRI showed high consistency across participants (**Fig. 2c-d**). To quantify the sources of variability in ISPC and PRI, we performed a variance decomposition across participants and frequencies for both rest and task episodes (see Methods). For ISPC, frequency-associated standard deviation was approximately 0.03 and participant-associated standard deviation was approximately 0.13. For PRI, frequency standard deviation was approximately 0.05 and participant standard deviation was approximately 0.23. These values were similar across episodes. Frequency-associated variability was small for both metrics, confirming that the phase-resolved connectivity patterns were stable across the frequency range examined. Participant-associated variability was larger, consistent with the spread of inter-subject standard deviations across electrode pairs (**Fig. 2c-d**, **Fig. S4**).

To test whether cortical distance accounted for the in-phase versus antiphase distinction, we grouped electrode pairs into four distance bins and separated them by connection type: homologous interhemispheric (8 pairs: FP1-FP2, F7-F8, F3-F4, T3-T4, C3-C4, T5-T6, P3-P4, O1-O2), non-homologous interhemispheric, and intrahemispheric (**Fig. 2e-f**). As distance increased, PRI shifted from values near 0 toward values near 1, indicating a transition from in-phase to antiphase clustering (**Fig. 2f**). Notably, however, homologous interhemispheric pairs did not follow this pattern. Despite spanning comparable distances to other pair types within the same bins (e.g., in the 110–142 mm range, mean distances were 125, 129, and 128 mm for homologous, non-homologous interhemispheric, and intrahemispheric pairs respectively; see **Table S1**), homologous pairs remained predominantly in-phase. Homologous connections traverse the corpus callosum, where axons are generally more heavily myelinated, resulting in shorter delays. This observation suggested that the transition from in-phase to antiphase was governed not by distance directly but by conduction delay.

The relationship between cortical distance and phase clustering is shown in **Fig. 2g**, where participant-median ISPC and PRI were plotted against distance in 10 mm bins. PRI increased as an approximate sigmoid with distance, rising from near 0 at short distances to approximately 1 at the longest distances, while ISPC dipped near the transition. This transition from in-phase to antiphase clustering, together with the accompanying dip in ISPC, motivated the modelling analyses that followed.

### A minimal model of delay-coupled populations reproduces phase clustering and the delay-driven transition

We next asked whether the observed phase clustering properties, including the transition from in-phase to antiphase with increasing cortical distance, could be explained by a minimal model of two delay-coupled excitatory neuronal populations. Since cortical distance covaries with conduction delay, and the homologous pair results suggested delay rather than distance as the controlling variable, we examined whether interpopulation delay alone could account for the transition.

The model consisted of two populations of leaky integrate-and-fire (LIF) neurons, each with intrapopulation coupling strength *G* and delay *d*, and interpopulation coupling strength *g* and delay *τ* (**Fig. 3a**). Simulations produced four qualitatively distinct regimes (**Fig. 3b–e**), which vary along two independent dimensions. The first is *synchronisation*, referring to within-population coherence: neurons fire in a coordinated, rhythmic manner, producing large-amplitude oscillations in population activity; in its absence, firing rates fluctuate irregularly around a mean. The second, defined only when both populations are synchronised, is *phase clustering*, referring to between-population phase agreement: the two populations may maintain a fixed phase difference (clustered) or their relative phase may drift over time (unclustered). Together these dimensions yield four regimes: unsynchronised irregular firing, synchronised but unclustered oscillations, in-phase clustered oscillations, and antiphase clustered oscillations.

**Fig. 3.**
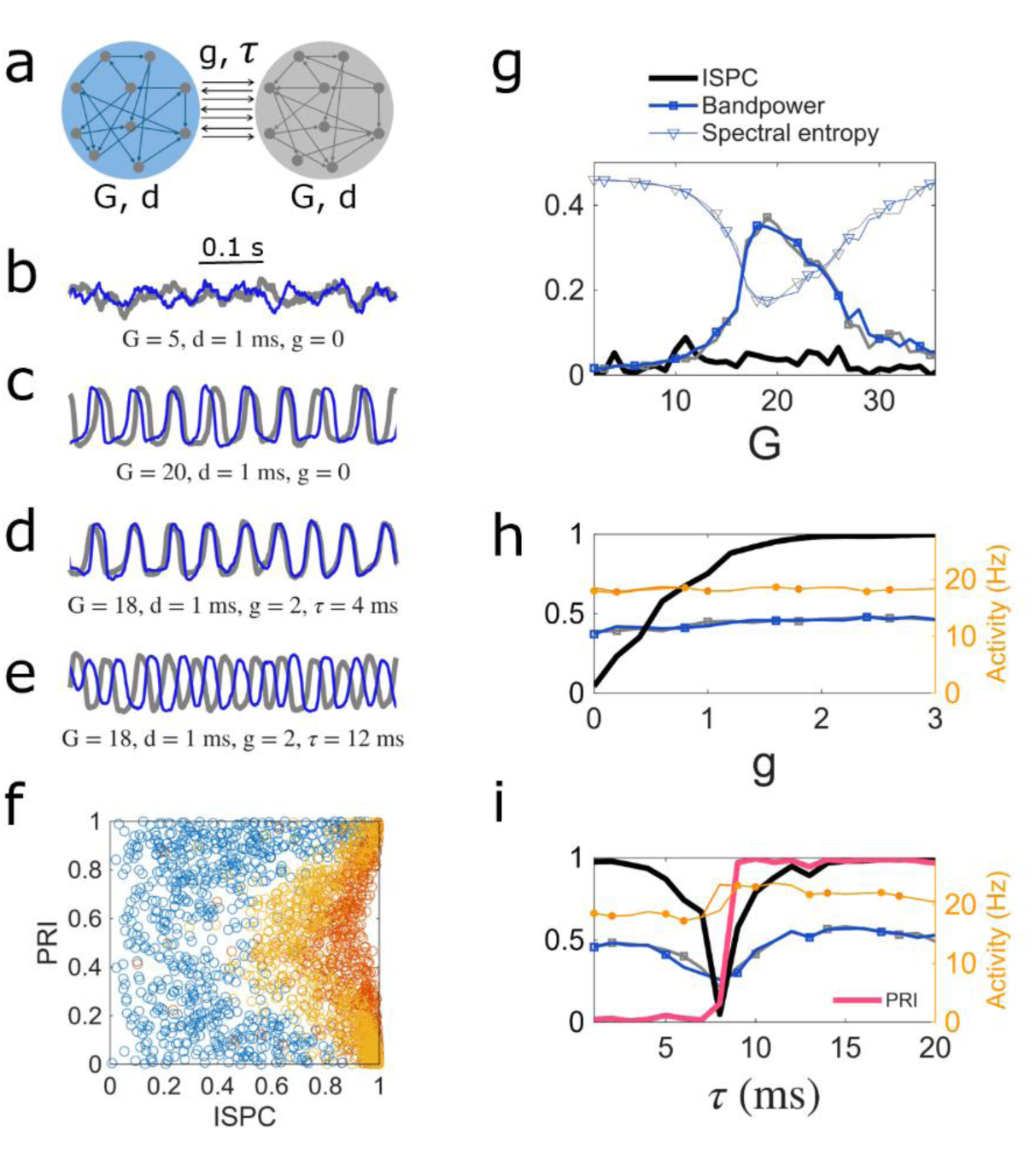
Delay-coupled excitatory populations exhibit a sharp transition from in-phase to antiphase clustering, governed by interpopulation delay. **a,** Model schematic. Two populations of excitatory neurons are coupled with intrapopulation connection strength *G* and delay *d*, and interpopulation strength *g* and delay *τ* . **b–e,** Representative dynamics from leaky integrate-and-fire (LIF) simulations (400 neurons per population), with parameters adjusted to produce four qualitatively distinct regimes: (**b**) unsynchronised, irregular firing; (**c**) synchronised, unclustered; (**d**) in-phase clustered; and (**e**) antiphase clustered. Traces show 0.5 s time segments of smoothed instantaneous population firing rates (spikes per neuron, 1 ms bins) for each population (blue, grey). **f,** ISPC and PRI computed from the simulated population activity time series, for each combination of coupling strength (*g*, 26 values uniformly selected from 0–1) and delay (*τ*, 20 values from 1–20 ms). Results are shown for time windows *W* = 1 s (red), 2 s (yellow), and 3 s (blue). **g,** Intrapopulation synchronisation in the absence of interpopulation coupling (*g* = 0). ISPC (black) and two measures of within-population synchronisation, relative band power near the peak frequency (squares) and spectral entropy (triangles), are shown for each population (blue, grey) as a function of intrapopulation connectivity *G*. Entropy values rescaled and shifted for visibility. External input: 1,200 spikes/s per neuron for both populations. **h,** Phase clustering emerges at weak interpopulation coupling. ISPC (black) increases sharply with *g*, while population firing rates (yellow and orange circles for the two populations) and synchronisation (blue and grey squares) remain approximately constant. Parameters: *G* = 20, *d* = 1 ms, *τ* = 1 ms. **i,** Interpopulation delay governs the transition from in-phase to antiphase clustering. ISPC (black) and PRI (red) are shown as a function of *τ*. ISPC dips near the transition, accompanied by a small increase in activities (circles) and a small dip in within-population synchronisation (squares). Parameters: *G* = 20, *d* = 1 ms, *g* = 2.

We computed ISPC and PRI for simulated population activities using parameters values chosen across a uniformly spaced grid of interpopulation coupling strength *g* and delay *τ*. The resulting distributions in the ISPC-PRI plane approximately reproduced the characteristic arc shape observed in the EEG data, with the arc sharpening as the analysis window *W* increased (**Fig. 3f**).

To isolate the contributions of intra- and interpopulation coupling, we first examined synchronisation in the absence of interpopulation connections (*g* = 0). As intrapopulation coupling *G* increased, each population transitioned from irregular firing to synchronised oscillation, marked by a dip in spectral entropy and a rise in band power near the peak frequency (**Fig. 3g**). With nonzero intrapopulation delay, synchronisation occurred within a bounded window of *G* values; however, when the intrapopulation delay was set to zero, there was a single threshold with no upper bound (**Fig. S5**). ISPC remained low throughout, confirming that interpopulation coupling was required for phase clustering to emerge.

With nonzero interpopulation coupling (*g* > 0 ), ISPC rose rapidly even at weak coupling strengths, while population firing rates and within-population synchronisation remained approximately constant (**Fig. 3h**). This demonstrated that phase clustering between populations could be established without substantially altering the dynamics within each population.

Finally, we examined the effect of interpopulation delay *τ* on phase clustering. As *τ* increased, PRI shifted abruptly from near 0 to near 1, indicating a transition from in-phase to antiphase clustering (**Fig. 3i**). Near the transition, ISPC dipped, mirroring features observed in the EEG distance-dependence data (**Fig. 2g**), and firing rates showed a small increase. This behaviour repeated approximately periodically with increasing *τ* (**Fig. S7**).

### Two competing instabilities govern in-phase and antiphase clustering

The LIF simulations demonstrated that interpopulation delay governed the transition from in-phase to antiphase clustering but left open the question of what mechanism underlies this transition. We therefore turned to a population density model amenable to analytical treatment, seeking to understand why ISPC dips and oscillation frequency jumps near the transition, and what determines the critical value of *τ* at which the transition occurs.

In the population density formulation, each population was described by a probability density *ρ*(*v*, *t*) or *ρ*′(*v*, *t*) representing the distribution of membrane potentials across neurons at time *t*. The two populations were coupled with the same structure as the LIF model: intrapopulation strength *G* and delay *d*, interpopulation strength *g* and delay *τ* (**Fig. 4a**). Given the same parameter values, the population density model reproduced the four dynamical regimes observed in the LIF simulations (**Fig. 4b-e**). One difference was that in the synchronised but unclustered regime, the deterministic population density model exhibited a fixed phase offset determined by initial conditions, whereas in the LIF simulations the phase offset drifted over time, producing low ISPC. **Fig. 4b-e** clearly show the populations settling on four different regimes despite starting from identical initial conditions.

**Fig. 4.**
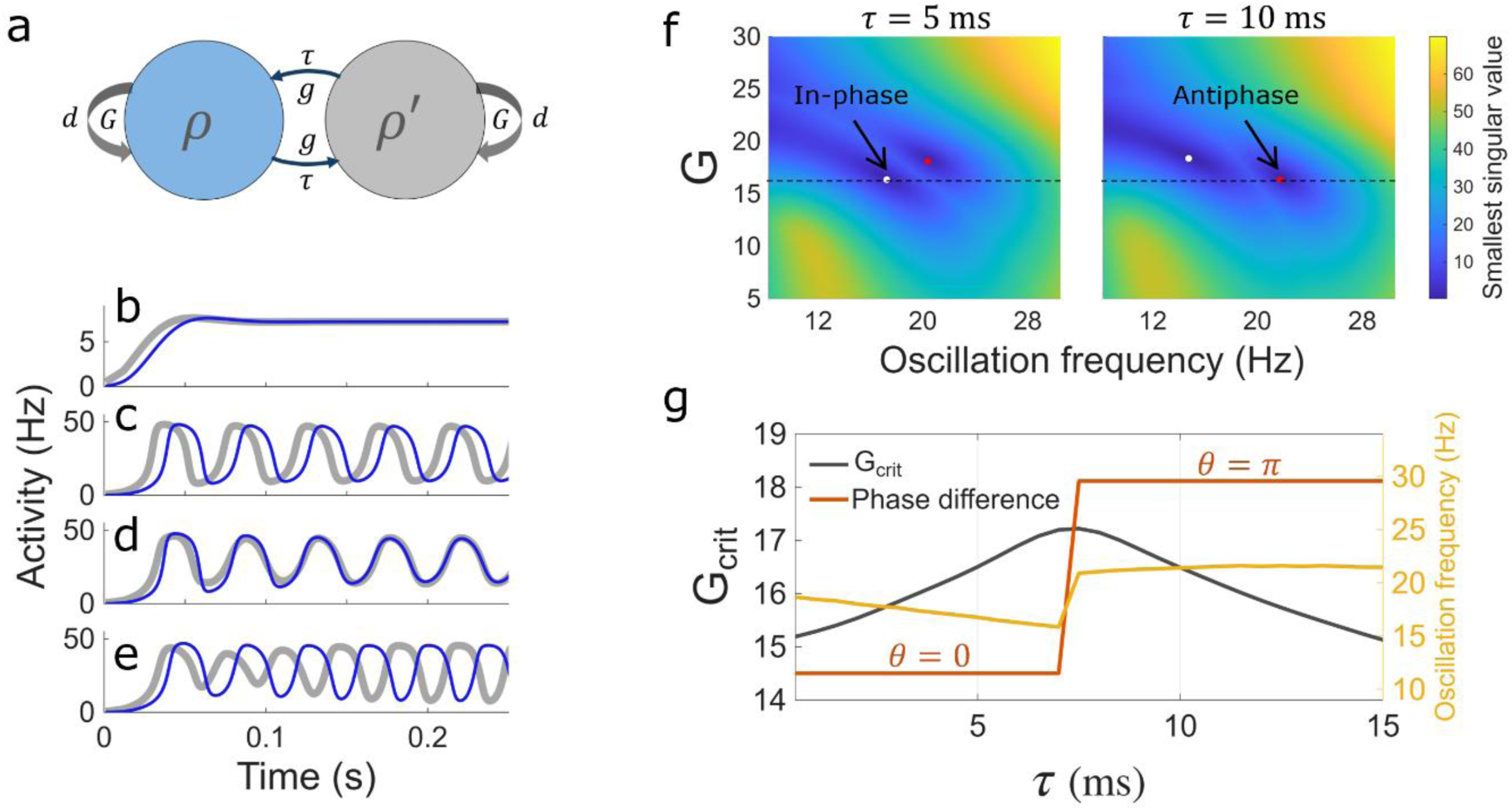
Linear stability analysis shows that in-phase and antiphase clustering arise from distinct Hopf bifurcations. **a,** Model schematic. Two populations, each described by a probability density *ρ* and *ρ*′, are coupled with intrapopulation strength *G* and delay *d*, and interpopulation strength *g* and delay *τ*. **b–e,** The population density model reproduces the four dynamical regimes observed in LIF simulations, given the same parameter values. (**b**) Unsynchronised: after initial transients, both population activities converge to the same constant value. (**c**) Synchronised but unclustered: each population oscillates, but with a phase offset determined by initial conditions; in the absence of noise, this offset remains fixed, whereas in LIF simulations it drifts, producing low ISPC over long time windows. (**d**) In-phase clustered: both populations rapidly converge to synchronous oscillation with zero phase lag. (**e**) Antiphase clustered: populations rapidly converge to oscillation with π phase lag. **f,** Linear stability analysis of the unsynchronised equilibrium. The smallest singular value of the stability matrix, i*ω* − M(i*ω*), is plotted as a function of intrapopulation coupling strength *G* and oscillation frequency *ω*/(2π). Two distinct foci appear where the singular value approaches zero, corresponding to in-phase and antiphase instabilities. As interpopulation delay *τ* increases, the foci rotate approximately clockwise around a common centre. The dominant instability (lowest *G_crit_*) swaps from the in-phase to the antiphase focus at a critical *τ*. Heatmaps are shown for *τ* = 5 ms (left, in-phase dominant) and *τ* = 10 ms (right, antiphase dominant). Parameters: *d* = 1 ms, *g* = 1, total input per population 1,580 Hz. **g,** Critical coupling strength *G*_crit_ (black, left axis) and oscillation frequency (yellow, right axis) as a function of *τ* . The phase difference between the two populations’ unstable modes (red) steps from 0 to π at the transition, as labelled on the plot.

At the crossover between in-phase and antiphase dominance, *G*_crit_ remains approximately constant while the oscillation frequency jumps abruptly.

We performed a linear stability analysis of the unsynchronised equilibrium to identify conditions under which the system would transition to oscillatory behaviour. Loss of stability, corresponding to emergence of sustained oscillations, occurred when the stability matrix became singular for some oscillation frequency *ω* > 0. We located these bifurcation points by computing the smallest singular value of the stability matrix across a grid of intrapopulation coupling strength *G* and oscillation frequency *ω*. This analysis revealed two distinct foci where the singular value approached zero, corresponding to two competing instabilities (**Fig. 4f**). At each focus, we extracted the associated eigenvector and computed the phase difference between the two populations’ perturbation modes: one focus corresponded to in-phase (*θ* = 0) and the other to antiphase (*θ* = π) oscillations.

As interpopulation delay *τ* increased, the two foci rotated approximately clockwise around a common centre in the *G*-*ω* plane (**Video S1**). For small *τ*, the in-phase focus had the lower critical coupling *G_crit_*, meaning the system first lost stability to in-phase oscillations as *G* increased. Beyond a transitional value of *τ*, the antiphase focus took over as the dominant instability (**Fig. 4f**, comparing heatmaps at *τ* = 5 ms and *τ* = 10 ms).

The behaviour near the transition is shown in **Fig. 4g**, with LIF simulations confirming the analytical predictions (**Fig. S6**). Before the transition to antiphase, as *τ* increased *G_crit_* rose slowly, making it progressively harder for oscillators to synchronise. After the transition, further increase in *τ* caused *G_crit_* to decrease, facilitating synchronisation. This explained a feature observed in the EEG data, namely an initial decrease in ISPC with increasing distance, followed by an increase in ISPC after the transition to antiphase (**Fig. 2g**). As *τ* increased through the transition, *G_crit_* remained approximately constant while the oscillation frequency jumped abruptly, and the phase difference stepped from 0 to *π*. This explained the frequency jump in the LIF simulations which arose because the in-phase and antiphase modes had different characteristic frequencies. The sharp transition was caused by the discrete swap in dominance between two distinct instabilities.

### Task-evoked changes in phase relationship are detected by PRI where ISPC shows little change

Having established that interpopulation delay governs the transition from in-phase to antiphase clustering, we next asked whether this distinction has functional relevance. Specifically, we examined whether task performance differentially affected in-phase and antiphase connections, and how ISPC and PRI captured task-evoked changes.

We computed the change in ISPC and PRI between the first task episode and the preceding rest episode (ΔISPC = T1 − R1; ΔPRI = T1 − R1) for each electrode pair and frequency using *W* = 32 s. Plotting these changes revealed that many electrode pairs showed substantial ΔPRI despite showing little or no ΔISPC (**Fig. 5a-b**). The pair with the largest shift toward in-phase (most negative ΔPRI) was C3–C4, the connection linking bilateral motor cortices, a finding consistent with the bimanual nature of the laparoscopic task.

**Fig. 5.**
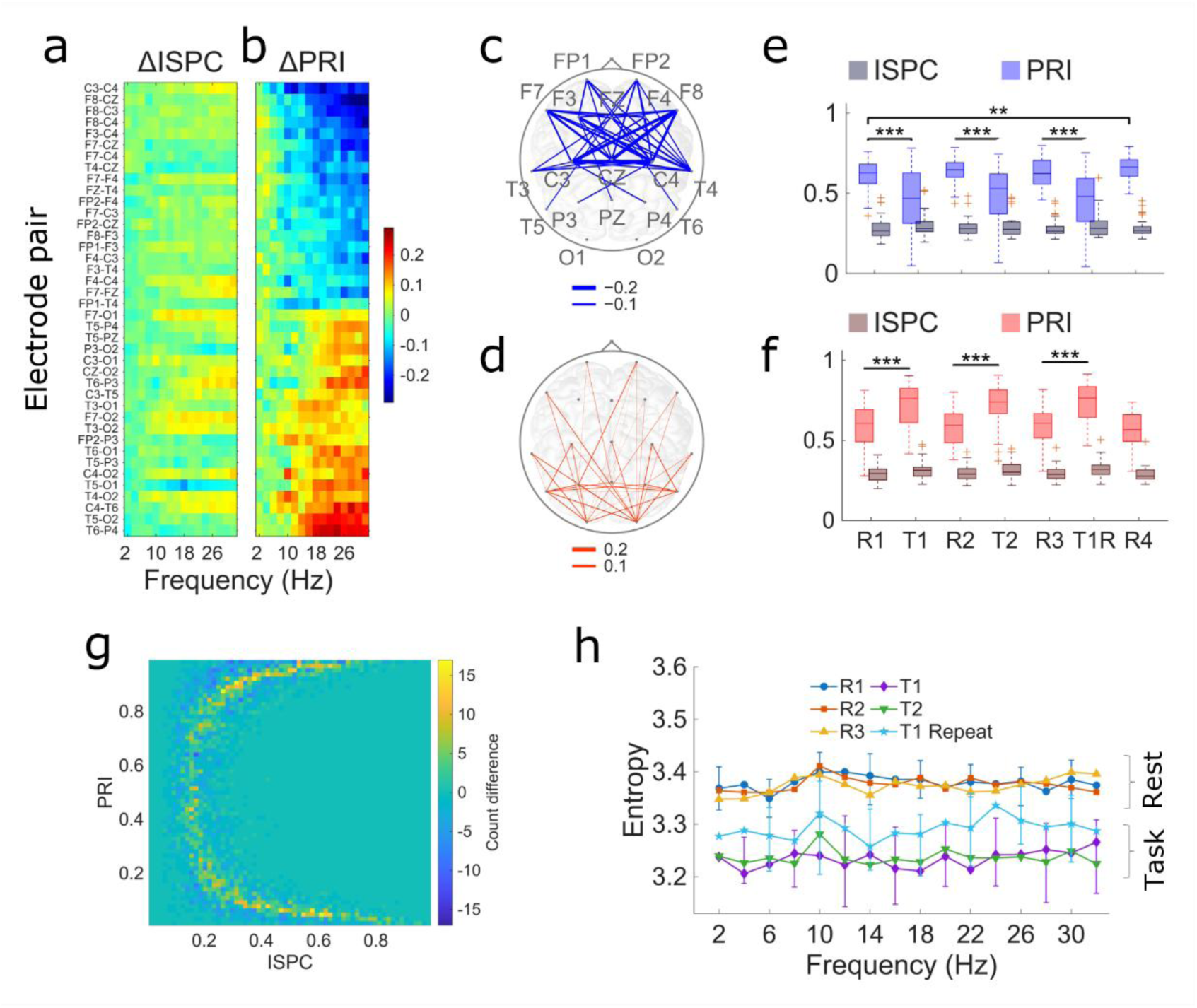
Task-related changes in phase synchrony characterised by ISPC and PRI. **a–b,** Change in ISPC (a) and PRI (b) between the first task episode and the first resting episode (ΔISPC = T1 − R1; ΔPRI = T1 − R1) for selected electrode pairs, plotted as a function of frequency. Pairs are sorted by ΔPRI in ascending order. For clarity, only the 20 pairs with the largest negative ΔPRI and the 20 pairs with the largest positive ΔPRI are shown. **c,** Topographic map of electrode pairs with negative ΔPRI, corresponding to connections that shifted toward in-phase synchrony during task performance. **d,** Topographic map of electrode pairs with positive ΔPRI, corresponding to connections that shifted toward antiphase synchrony during task. **e,** PRI (blue) and ISPC (grey) as a function of episode for the network identified in (c), with boxes indicating the interquartile range (25th–75th percentile) across participants. Significant differences between episodes are indicated (** *p*<0.01, *** *p*<0.001). **f,** As in (e), for the network identified in (d). **g,** Difference heatmap in the ISPC–PRI plane (T1 − R1), for all electrode pairs and participants, illustrating a narrowing of the arc-shaped distribution during task relative to rest. **h,** Entropy of the joint ISPC–PRI distribution as a function of frequency for each episode, showing consistently lower entropy during task episodes relative to rest episodes, consistent with the tightening of the arc observed in (g).

To identify task-modulated networks, we selected the electrode pairs with the largest negative and positive ΔPRI values. Pairs shifting toward in-phase connectivity formed a distributed network spanning frontal, central, and parietal regions (**Fig. 5c**), while pairs shifting toward antiphase formed a distinct network (**Fig. 5d**).

We then tracked ISPC and PRI across all experimental episodes for these two networks. For the networks that shifted toward in-phase (**Fig. 5e**) or antiphase (**Fig. 5f**), PRI showed significant changes during task episodes relative to rest, while ISPC showed smaller and less consistent changes. These results indicated that task performance modulated phase relationships in ways that would be obscured by examining ISPC alone.

Beyond changes in individual electrode pairs, we observed that the overall shape of the joint ISPC-PRI distribution changed between rest and task. The arc-shaped distribution that characterised resting connectivity became narrower during task performance, as visualised in the difference heatmap (**Fig. 5g**). To quantify this tightening, we computed the Shannon entropy of the joint ISPC-PRI distribution for each episode and frequency. Entropy was consistently lower during task episodes than rest episodes across all frequencies examined (**Fig. 5h**). A phenomenological stochastic model of phase difference dynamics replicated this tightening effect. Increasing the mean-reversion speed toward preferred phase differences, while holding stationary variance constant, produced a narrower arc in the ISPC-PRI plane (**Fig. S8**). Notably, the repeat task episode (T1R) showed entropy values intermediate between the initial task episodes (T1, T2) and the rest episodes.

Together, these results demonstrated that the phase relationship captured by PRI was sensitive to cognitive state, suggesting that PRI used alongside ISPC may provide a more complete characterisation of functional connectivity and its modulation by task demands, with potential applications as a biomarker of cognitive engagement.

## Discussion

We observed a distance-dependent transition from in-phase to antiphase connectivity in EEG and interpreted this as reflecting conduction delay rather than distance per se. A similar distance-dependent transition has previously been reported without, however, making a link to conduction delay^30^. Linking delay to the transition was supported by the finding that homologous interhemispheric pairs remained in-phase despite spanning long distances, consistent with faster callosal conduction. A model of delay-coupled excitatory populations confirmed this interpretation, reproducing the transition and revealing its mechanistic basis in competing instabilities. Our finding that task performance modulates phase relationships in a frontoparietal network suggested that this organisation was not merely structural but dynamically engaged during cognitive demands. The observed ISPC-PRI distribution was also evident in preliminary results from independent datasets^34–36^ (**Fig. S9a-c**), suggesting that the bimodal phase organisation generalises across populations and recording contexts. These findings suggest that interareal connectivity is organised primarily by phase relationship as a dynamical state variable, with transitions between discrete in-phase and antiphase regimes rather than continuous variation in coupling strength, a structure obscured by metrics that collapse phase information or exclude zero-lag components.

### Prior evidence for bimodal phase relationships

The bimodal distribution of phase relationships, centred near 0 and π, is consistent with prior reports from intracortical and intracranial recordings. Near-zero phase lag has been found in theta synchronisation between amygdala and hippocampus^37,38^, gamma synchronisation between rhinal cortex and hippocampus^9^ and between visual areas V1 and V2^39^, long-range beta synchronisation across visual, parietal and motor cortices in cats^40^, and gamma synchronisation across neocortical regions in humans^41^. Bimodal distributions centred near 0° and 180° were observed in monkey prefrontal and posterior parietal cortex, with rapid task-dependent transitions between modes^42^. Intracranial EEG showed that in-phase interactions were highly specific to homotopic regions, ruling out volume conduction^25^. Furthermore, source-level analysis of high-density EEG confirmed pervasive zero-lag interactions and showed that metrics excluding zero-lag penalise prediction of age and cognition^31^.

Several mechanisms have been proposed to account for zero-lag synchronisation despite conduction delays: common input, mutual excitatory-inhibitory dynamics^43^, spike doublets in interneurons^44,45^, thalamic relay^46,47^, or emergent network effects^4^. Our findings are consistent with a network effect, as delayed mutual excitation between two populations suffices to produce both zero-lag and π-lag synchronisation.

### The transition reflects delay-governed dynamics

Our observation that homologous interhemispheric pairs remain in-phase despite spanning long distances (**Fig. 2**e-f; **Table S1**) supports the interpretation that conduction delay, rather than distance per se, governs the transition. Homologous connections traverse the corpus callosum, where axons are more heavily myelinated, resulting in faster conduction. The transition from in-phase to antiphase bears the hallmarks of the phase-flip bifurcation described in delay-coupled dynamical systems^32,48^. This transition is typically accompanied by amplitude death, a transient suppression of oscillations, presumably responsible for the ISPC dip we observed (**Fig. 2**g). The phase-flip bifurcation has been demonstrated in physical systems including coupled candle-flame oscillators^33^ and joint hand-clapping^49^. Our stability analysis provides an account of this phenomenon in a neural population model, showing that two distinct instabilities exchange dominance as delay increases (**Fig. 4**f-g).

### A minimal model captures the empirical phenomenology

Our model consists of two excitatory populations coupled with delay, with no inhibition and no complex network topology. We analysed it through two complementary approaches: direct simulations of leaky integrate-and-fire neurons receiving stochastic input, and a deterministic population density formulation amenable to stability analysis. Despite its simplicity, the model captures multiple features of the empirical data: the arc-shaped joint distribution in the ISPC-PRI plane (**Fig. 3**f), bimodal clustering at in-phase and antiphase, the dip in ISPC near the transition (**Fig. 3**i), and the sharp delay-governed transition. These features emerged without parameter tuning.

The success of such a minimal model suggests that in-phase and antiphase organisation reflect a fundamental dynamical phenomenon that cortical networks may exploit for functional purposes. In addition, the finding that weak interpopulation coupling (*g* ≪ *G*) suffices for long-range phase clustering is consistent with small-world architecture^50^, known to characterise cortical connectivity^51^. Modulating *g* may provide an energetically efficient mechanism for transiently linking distant areas^1^.

### Phase differences evolve as noisy attraction to preferred states

The arc-shaped distribution in the ISPC-PRI plane was replicated by a stochastic Ornstein-Uhlenbeck model of phase difference dynamics (**Fig. S8**). Each electrode pair maintains a noisy mean-reverting phase relationship with a preferred phase difference that varies across pairs. The model reproduced the arc only when mean-reversion was strongest near 0 and π, suggesting these are attractors. The model also replicated task-evoked tightening of the arc (**Fig. 5**g-h): increasing mean-reversion speed while holding stationary variance constant produced a narrower arc (**Fig. S8b**), suggesting that attraction to preferred phase differences may be modulated by task demands, potentially through changes in cortical excitability. This phenomenological account complements the mechanistic model by describing how phase relationships evolve stochastically around preferred values.

### Volume conduction cannot explain the observed patterns

Several lines of evidence argue against volume conduction. First, antiphase connectivity is unlikely to arise from volume conduction, yet it constitutes a substantial portion of our findings. Second, the distance-dependent transition from in-phase to antiphase matches predictions from conduction delay, not the monotonic decay expected from passive spread. Third, task-evoked modulation implies neural dynamics; it is unclear how volume conduction could selectively increase for a specific frontoparietal network during task.

Bimodal phase distributions were found in monkey prefrontal and posterior parietal cortex, where antiphase correlations and task-dependent dynamics ruled out volume conduction^42^. In addition, zero-lag synchrony was present in human intracranial recordings where volume conduction drops almost completely within short distances (<20 mm)^25^. Since long-distance zero-lag connectivity exists intracranially, it cannot be automatically dismissed from scalp EEG. In our study, we applied surface Laplacian transformation prior to analysis, attenuating the broad spatial components associated with volume conduction. Without this filtering, nearly all electrode pairs clustered at PRI near 0 with high ISPC; following filtering, the full arc was recovered (**Fig. S9**).

### PRI reveals task-related dynamics invisible to ISPC

ISPC quantifies clustering strength independent of the underlying phase relationship; PRI captures this complementary dimension, ranging from 0 (in-phase) to 1 (antiphase). Task-evoked changes in phase relationships were detected by PRI in electrode pairs where ISPC showed little or no change (**Fig. 5**a-b). The pair with the largest shift toward in-phase during task was C3-C4, linking bilateral motor cortices, consistent with the bimanual nature of the laparoscopic task (**Fig. 5**c). Networks identified by PRI changes showed significant modulation across experimental episodes, while ISPC changes were smaller and less consistent (**Fig. 5e-f**).

### Task demands shift frontoparietal connectivity toward in-phase

Task performance shifted a distributed frontoparietal network toward in-phase connectivity (**Fig. 5**c), with PRI decreasing significantly during task relative to rest (**Fig. 5**e). A separate network shifted toward antiphase (**Fig. 5**d, **Fig. 5**f). The arc narrowed during task (**Fig. 5**g), quantified by reduced Shannon entropy of the joint distribution across all frequencies (**Fig. 5**h). Intermediate entropy during the repeat task episode (T1R) may reflect a transition toward automated performance, with less active coordination than initial learning yet more than rest.

These findings align with intracortical recordings in monkeys where frontoparietal correlations increased during working memory delay periods and relative phase could shift by up to 180° following error trials^42^.

### Limitations

Several limitations should be noted. The sample size was modest (*N* = 31), drawn from a single dataset and analysis was conducted at sensor level with 19 electrodes, limiting spatial resolution and anatomical interpretation. These could be addressed by dense arrays with source reconstruction. In addition, resting epochs were short (2 minutes) and participants stood during task, possibly introducing movement artefacts.

The model lacks inhibition, known to shape oscillatory dynamics and it uses a single fixed delay, whereas distributed delays^52^ may better capture variability in conduction times; and anatomical or diffusion tensor imaging-based estimates of delay^53^ could sharpen model-data correspondence. Furthermore, our stability analysis identified when the equilibrium loses stability; however, it did not formally prove the resulting oscillations are attractors, although simulations strongly suggest they are.

Finally, the observation that the transition from in-phase to antiphase occurs at roughly the same cortical distance across all frequencies presents interpretive challenges. For delay-coupled oscillators, this transition is expected at a delay of approximately one quarter of the oscillation period. For shorter delays, spikes from one population arrive when the other is in its excitable phase, reinforcing in-phase oscillation; for delays between one quarter and three quarters of the period, spike arrivals sustain antiphase oscillation (**Fig. S10**). Our simulations confirm that when delay falls within these ranges, phase differences are attracted toward 0 or *π* (**Fig. 3**i). For example, at 10 Hz (period 100 ms), in-phase clustering is expected for delays below approximately 25 ms. At the observed transition distance of approximately 100 mm (**Fig. 2g**), this corresponds to conduction velocities above 4 m/s, which is within the physiological range. If the transition distance scaled with period, connections with double the frequency at 20 Hz would have half the transition distance at approximately 50 mm, keeping velocity unchanged. Yet our data show that transition distances remain similar across the examined frequency range, implying, according to the above reasoning, that velocity must scale with frequency. Since conduction velocity is determined by axonal properties rather than oscillation frequency, this implication is physiologically implausible. This puzzle may reflect the broadband nature of local cortical activity, where the mapping between narrowband-filtered EEG and underlying population dynamics remains to be clarified.

### Predictions

Dual-site transcranial alternating current stimulation (tACS) with controlled phase offsets could test whether in-phase or antiphase stimulation more effectively modulates connectivity between cortical sites. Previous work showed that connectivity changes after dual-site tACS depend on phase lag between sites, with effects further modulated by conduction delays^54^. Our model predicts that the optimal phase relationship for entrainment should depend on effective delay, with in-phase stimulation more effective at short delays.

## Methods

The experimental design and data collection have been described in detail previously^55,56^. The following summarises the key aspects relevant to the present analysis.

### Participants

A total of 38 healthy adult volunteers without prior experience in laparoscopic surgery participated in the study. One participant could not be fitted with the EEG cap, four were excluded due to technical problems, and two were excluded due to timestamp recording issues, leaving a final sample of N = 31 participants (17 females, 14 males, mean age 21.61 ± 2.12 years). All participants provided written informed consent prior to the study and received gift vouchers for their participation. The study was approved by the Ethical Committee of the College of Science and Technology at Nottingham Trent University, and all methods were performed in accordance with the relevant guidelines and regulations. Representative participants shown in **Fig. S9** were drawn from previously published datasets^34–36^ collected under separate institutional ethics approvals that permitted secondary analysis for research purposes.

### Experimental design

Participants performed two laparoscopic training tasks of comparable difficulty, Ring Transfer and Threading, on a laparoscopic simulator, with the first task repeated near the end of the session to assess performance gains. The experimental sequence consisted of an initial reaction task (T0), an initial rest episode (R1), the first training task (T1) selected randomly from the two task types, a rest episode (R2), the second task (T2), another rest (R3), a repeat performance of the first task (T1R), and a final rest episode (R4), each rest lasting 2 minutes. EEG was recorded throughout from 19 channels at standard 10–20 sites. Performance was measured as task completion time, with a maximum allowed time of 15 minutes, and errors were tracked for the Ring Transfer task. Subjective cognitive load was assessed after each task using the NASA-TLX questionnaire. Participants stood in front of the laparoscopic simulator box during task performance. The Ring Transfer task involved grasping, lifting, and relocating rings between four rods (A to D) using both surgical instruments, following a prescribed sequence of transfers. The Threading task consisted of passing a piece of string through seven labelled holes using both surgical tools and both hands. An auditory secondary task was performed concurrently throughout the experiment, in which participants responded to a series of beeps as quickly as possible by pressing a foot pedal, simulating the distractions that may arise in a realistic surgical setting.

### EEG pre-processing

Raw EEG data underwent a series of pre-processing steps to remove non-brain signals. Segments containing high-amplitude artifacts likely originating from gross body movements were identified and removed using a 1 s sliding window. Electrodes with kurtosis values exceeding 5 were removed as invalid channels. The signals were band-pass filtered between 0.16 Hz and 40 Hz to reduce slow drifts and high-frequency artifacts, then down sampled to 200 Hz. Independent Component Analysis (ICA) decomposition using the Extended-Infomax algorithm was applied to the filtered data to segregate artefactual components such as eye blinks and movements. The ADJUST method was used to automatically detect and remove artifact-related components, supplemented by visual inspection of the power spectrum, time series, and topographic maps of each component. The pre-processed data were then reconstructed for further analysis.

### Inter-Site Phase Clustering (ISPC)

Inter-Site Phase Clustering (ISPC), also known as Phase Locking Value (PLV), was calculated for each unordered pair of electrodes within 1 Hz wide frequency bands centred at 1, 2, …, 40 Hz. ISPC measures the consistency of the inter-site phase difference over time but does not convey information about the nature of that difference itself. To calculate ISPC, narrow-band analysis was used to avoid potential loss of information from pooling into traditional wider frequency bands and to generate more reliable phase estimates^57^. To minimise distortion by volume conduction, a Laplacian spatial filter was applied, subtracting from each electrode’s instantaneous signal the mean of its nearest neighbours to estimate the local current source density^58^. For each electrode pair, the associated cortical distance was computed from the MNI coordinates of the primary neuronal populations underlying each electrode^59^.

For each frequency band the pre-processed signals were first narrowband filtered, and the Hilbert transform applied to extract the instantaneous phase of each signal. For each pair of electrodes, the complex-valued instantaneous phase clustering vector for the *n*th time window was calculated as 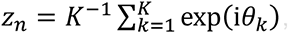, over a window of size *W* containing *K* samples (**Fig. S2a**). Examples of phase difference vectors are illustrated as dots on the complex plane in **Fig. S1** and **Fig. 1a-c**. ISPC was then calculated as *I_n_* = |*z_n_*|. To verify the robustness of our phase extraction calculations, we also applied a zero-crossing method as an alternative to the Hilbert transform. Comparing the two approaches revealed only minor differences in the results **Fig. S3**.

### Phase Relationship Index (PRI)

ISPC measures the strength of phase clustering but does not convey information about the nature of the phase relationship itself. To address this, we introduced the Phase Relationship Index (PRI), defined as *P_n_* = |*θ_n_*|/π, where *θ_n_* = Arg(*z_n_*) . PRI ranges from 0 (in-phase synchrony) to 1 (antiphase), and can capture any phase relationship, not exclusively in- or antiphase. The absolute value of *θ_n_* was used for several reasons. First, without it PRI would range over (−1, 1), where the extreme values −1 and 1 would both represent antiphase synchrony but appear numerically and visually distinct, which would be misleading. Second, and more importantly, averaging signed values across subjects, frequencies, or channel pairs might cause positive and negative antiphase contributions to cancel, giving the erroneous impression of in-phase clustering. Taking the absolute value ensures that PRI aggregates correctly and intuitively across all dimensions. Consequently, PRI does not indicate which electrode in a pair leads in phase; if directionality were of interest, the signed angle could be analysed directly.

### Mitigating volume conduction by using spatial filtering

A potential concern in EEG connectivity analysis is the contribution of volume conduction, which causes the instantaneous spreading of electrical potentials across the scalp and produces spurious zero-lag (in-phase) synchrony between electrode pairs. In the context of PRI, unmitigated volume conduction would bias measurements toward PRI ≈ 0 regardless of the true underlying phase relationship, inflating the apparent prevalence of in-phase connectivity. To address this, a surface Laplacian (current source density) spatial filter was applied to all EEG signals prior to analysis. The Laplacian is a well-established spatial high-pass filter that attenuates the broad, spatially smooth components associated with volume conduction while preserving local cortical sources^60^. To illustrate its effect, we computed ISPC-PRI distributions for a representative subject from an independent resting-state EEG dataset, with and without Laplacian filtering. Without filtering, nearly all channel pairs clustered at PRI ≈ 0 with high ISPC, consistent with volume-conduction-dominated in-phase clustering. Following Laplacian filtering, the full arc from in-phase to antiphase clustering (e.g. **Fig. 1**d) was recovered across channel pairs (**Fig. S9**).

### Entropy of the joint ISPC-PRI distribution

To quantify the dispersion of the joint distribution of ISPC and PRI, we computed its Shannon entropy. The ISPC and PRI values were binned into a two-dimensional histogram with bin indices *j* and *k*, yielding a normalised joint probability distribution *p_jk_* . The entropy was calculated as *H* = − ∑ *p_jk_ log*_2_(*p_jk_* ), where the sum was taken over all occupied bins. Lower entropy indicated a more concentrated distribution, corresponding to tighter clustering in the ISPC-PRI plane.

### Statistical validation of phase clustering direction

To statistically test the observation that phase relationships tend toward in-phase or antiphase, we applied a circular statistical analysis. The complex-valued phase clustering vector *z*(*e*, *s*, *f*, *p*; *W*) was obtained by averaging the instantaneous phase clustering vector *z_n_* across time windows within each episode. The phase angle *θ* = Arg(*z*) and the ISPC = |*z*| were then extracted for each combination of episode, subject, frequency, and channel pair. To test whether *θ* congregates around either 0 or *π*, a doubling transformation *θ*′ = 2*θ mod*(2*π*) was applied, which maps both 0 and *π* to the same point on the unit circle. The V-test^61^ for circular uniformity against a specified mean direction of 0 was then applied to *θ*′ pooled across subjects, yielding one p-value per frequency-electrode pair combination. Multiple comparisons were controlled using the Benjamini–Hochberg false discovery rate procedure at *α* = 0.05.

### Variance decomposition of ISPC and PRI across subjects and frequencies

For each metric (ISPC and PRI) we quantified the contributions of subjects and frequency bins to overall variability using a marginal-variance decomposition on the full 31 × 16 × 171 data array. For each electrode pair, ISPC/PRI values were arranged as subjects × frequencies; we computed (1) the subject-associated variance by first averaging across frequencies for each subject and then taking the variance of those subject means, and (2) the frequency-associated variance by first averaging across subjects for each frequency and then taking the variance of those frequency means. Variance estimates were averaged across the 171 electrode pairs to produce mean variance components for subjects and for frequencies for each episode. Ratios of subject-to-frequency variances were computed to summarise the relative magnitude of the two variance sources. All computations were performed in MATLAB (MathWorks), and variances are reported on the original metric of ISPC and PRI.

### Direct simulations

To examine the clustering dynamics observed in EEG data, we performed simulations of two coupled excitatory Leaky Integrate-and-Fire (LIF) neuron populations. Each neuron’s membrane potential evolved according to:

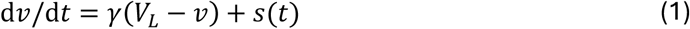

with the reset condition that when *v* reached the firing threshold *V_T_*_ℎ_, it was immediately reset to the resting potential *V_R_*. Each population contained *N* = 400 neurons, with initial membrane potentials drawn arbitrarily from the interval (*V_R_*, *V_T_*_ℎ_) . Connectivity was defined at the population level; each neuron projected on average to *G* post-synaptic targets randomly selected within its own population (intrapopulation connectivity), with spikes arriving after a conduction delay *d*. It additionally projected to an average of *g* neurons in the other population (interpopulation connectivity), with spikes arriving after delay *τ*. Each neuron also received Poisson-distributed external input at the constant rate *σ*⁰, representing drive from regions outside the simulated pair of populations. Neurons were pulse-coupled, meaning each arriving spike produced an instantaneous jump in the receiving neuron’s membrane potential: *v* → *v* + ℎ. This captured the essential effect of a synaptic conductance change occurring on a timescale fast relative to the membrane dynamics. The stimulus term accordingly took the form: *s*(*t*) = ℎ ∑ *δ*(*t* − *t_k_*) where *t_k_* denoted the arrival time of the *k*th spike. Parameter values *V_L_* = −65 *mV* (leakage potential), *V_T_*_ℎ_ = −35 *mV*, *V_R_* = −65 *mV*, and ℎ = 1 mV were chosen to reflect the characteristic properties of cortical pyramidal neurons^62^. Numerical integration was performed using the forward Euler method with a time step of *Δt* = 0.1 ms.

Choice of smaller time steps did not affect the results. The activity of a population (quantified as the firing rate, i.e. the number of spikes per neuron within adjacent 1 ms bins) was taken as a proxy for the EEG signal. To verify that results were not sensitive to this choice, simulations were also analysed using the total synaptic arrival rate per neuron as an alternative measure of population activity, which did not change the phase clustering results.

### Analysis of direct simulation results

The population activity resulting from direct simulations was used to quantify synchronization and phase clustering. When a population’s behaviour was oscillatory with a frequency *f_max_*, the power spectrum of its activity (or firing rate) was peaked around *f_max_*. Thus, we computed Frequency Band Power (FBP) by first determining, *f_max_*, and then determining the relative band power within a Δ*f* = 5 Hz wide band centred around *f_max_* . We also computed the spectral entropy as *S* = − ∑ *p_k_ log*_2_(*p_k_*) where *p_k_* was the spectral power at the *k*th frequency. For clustering, ISPC and PRI were computed from the activities of the pair of simulated populations.

### Population model

Excitatory neurons that dynamically follow the LIF model were characterised at population level by *ρ*(*v*, *t*), the probability that a member of the population is at membrane potential *v* at time *t*. Under general conditions it has been shown that^63–65^

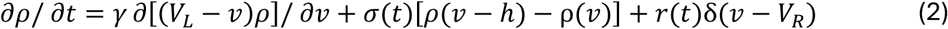

where the first term on the right represents leakage, the second term represents the changes of state due to arrivals of synaptic input at the rate *σ*(*t*) per neuron, and the last term with the Dirac delta function arises from neurons that fired being reinstated at *v* = *V_R_* . The latter occurs at the rate *r*(*t*), the firing rate per neuron. We used (2) to model the dynamics of *ρ* with boundary conditions *ρ*(*V_T_*_ℎ_, *t*) = *ρ*(*v* → −∞, *t*) = 0 . As *ρ* was a probability, it satisfied 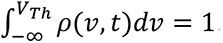.

The momentary fraction of *susceptible* neurons, i.e. those that fire immediately on receiving a synaptic input, was

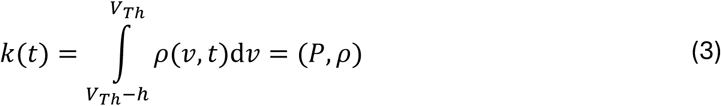

where (*P*, *ρ*) denoted the weighted integral of *ρ* over (−∞, *V_T_*_ℎ_), with kernel *P*(*v*) equal to unity on ( *V_T_*_ℎ_ − ℎ, *V_T_*_ℎ_ ) and zero elsewhere. This notation was selected as it allows some modifications to the synaptic dynamics to be accommodated through a change in *P*(*v*), leaving (*P*, *ρ*) unchanged. The activity or the firing rate of the population was *r*(*t*) = *σ*(*t*)*k*(*t*). We also introduced a time delay operator such that *T_d_ f*(*t*) = *f*(*t* − *d*) for any measurable function *f*(*t*), with *d* as the axonal conduction delay. The total synaptic input to a neuron thus equalled the external input *σ*^0^ plus the population’s own recurrent activity amplified by *G*, arriving with a delay *d*:

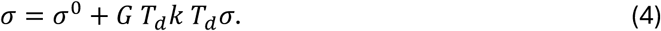

We next introduced dynamical operators, for leakage *Q^L^ρ* = *γ∂*[(*V_L_* − *v*)*ρ*]/*∂v*, and for excitatory input *Q^E^ρ* = *ρ*(*v* − ℎ) − *ρ*(*v*) + (*P*, *ρ*)*δ*(*v* − *V_R_*), to write (2) compactly as

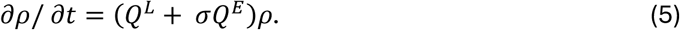

Henceforth we preferred to work with this form as it could accommodate, within a unified notation, potential modifications to the underlying neuronal dynamics, and admit future extensions, e.g. inhibitory interactions. Combining (5) with (4) provided the complete nonlinear dynamics of a single population with feedback gain *G* and delay *d*, and external input *σ*^0^:

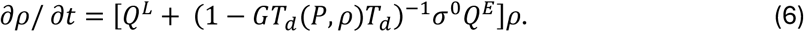

With constant external input and sufficiently weak connectivity, the population reached a steady equilibrium *ρ*_0_(*v*) that satisfied

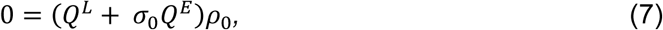

where *σ*_0_ = *σ*^0^ + *G k*_0_*σ*_0_ and *k*_0_ = (*P*, *ρ*_0_). The delay operator drops out of the equilibrium variables as they are constant. For simulations and stability analysis, (5) was discretised through a probability conserving numerical scheme that was 2^nd^ order in *v* and in time^65^.

### Coupled populations

We extended this approach to a pair of interacting populations to analyse the stability of its equilibria. For the sake of a concise analysis, it was assumed that the populations were configured symmetrically, i.e. they had the same size, membrane and synaptic dynamics and external inputs. The interpopulation connectivity strength was denoted *g* (per neuron average number of postsynaptic target neurons in the other population) with an interpopulation axonal conduction delay *τ*. The total input rate (4) was now extended to include an additional term representing the contribution from the other population:

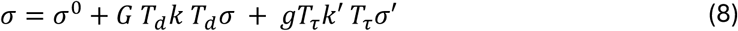

where, for the other population, the total input rate was *σ*′(*t*) and the susceptible fraction *k*′(*t*) = (*P*, *ρ*’), with the probability *ρ*’(*v*, *t*) representing its momentary state. While the first population was described by (5) and (8), the second population was described by

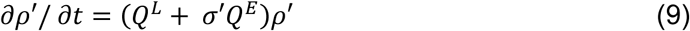

and

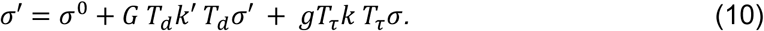

Due to the symmetry of the setup, the two populations’ equilibria were the same, *ρ*′_0_ = *ρ*_0_.

### Stability analysis

We introduced small perturbations about equilibrium so that

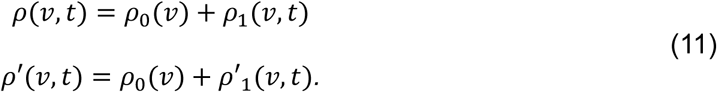

Linearised dynamics near equilibria were obtained by inserting (11) into (5) and (9) and keeping only terms first order in the perturbation variables *ρ*_1_ and *ρ*′_1_. The result was, for the first population,

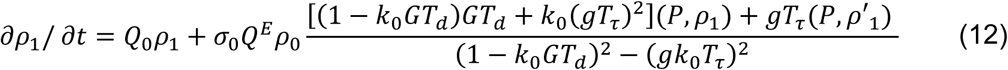

and, for the other population, an identical equation but *ρ*_1_ exchanged with *ρ*′_1_, where we have defined *Q*_0_ = *Q^L^* + *σ*_0_*Q^E^*.

For exponential solutions in the time domain, *ρ*_1_(*v*, *t*) = *u*(*v*)e*^μt^* and *ρ*′_1_(*v*, *t*) = *u*′(*v*)e*^μt^*, it followed that

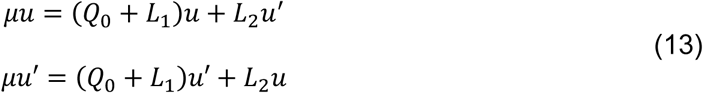

with

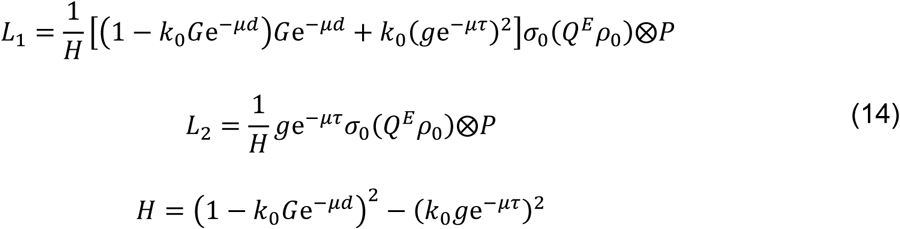

where the only terms that depend on *v* are (*Q^E^ρ*_0_) and *P*, and ⊗ denotes the outer product defined by (*a*⊗*b*)*c* = (*b*, *c*)*a* for any functions *a*(*v*), *b*(*v*), *c*(*v*).

For the case of uncoupled populations, i.e. *g* = 0, (13) reduces to two identical expressions equivalent to a previously published single population stability analysis^63^.

Discretising the operators allowed us to write the functions *u* and *u*’ as column vectors **u** and **u**′, which we concatenated into a single vector **φ**. Then (13) could be written *μ***φ** = M**φ** with

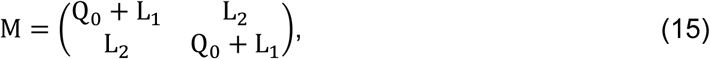

using the numerical approximations to the dynamical operators in (13)-(14). To locate neutral stability curves in parameter space we forced *μ* to be pure imaginary, *μ* = i*ω*, implying that (i*ω* − M)**φ** = 0. Then, we performed grid search on (*G*, *ω*) to determine the pairs (*G*_c*rit*_, *ω*_c*rit*_) for which the stability matrix i*ω* − M was singular, i.e. admitted a non-trivial null vector **φ*_c_***. Since the smallest singular value *σ_min_*(i*ω* − M) vanishes if and only if i*ω* − M is singular, neutral stability was identified as points where *σ_min_* = 0 . This approach was numerically preferable to directly evaluating det(i*ω* − M ) as singular value decomposition (unlike calculating the determinant) remains well-conditioned for large matrices.

From **φ*_c_*** we extracted **u** and **u**′, and estimated their phase difference as 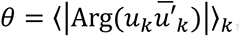, i.e. the absolute wrapped phase difference averaged across elements *k* . The search for neutral stability revealed that *σ_min_* = 0 (to numerical precision) with *ω_crit_* > 0 (oscillatory instability) occurred at two distinct foci, one with *θ* = 0 (in-phase) and the other *θ* = π (antiphase). Since the system first loses stability at the focus with the lower critical coupling *G_crit_*, we identified this as the relevant bifurcation point, with the character of the emerging oscillations determined by its associated *θ*. We note that the condition *σ_min_* = 0 was also met along *ω* = 0, corresponding to a steady-state bifurcation of the equilibrium rather than an oscillatory instability; these points were excluded from the analysis. The structure of the instability, with a pure imaginary eigenvalue and oscillatory solutions emerging beyond *G*_crit_, was consistent with a Hopf bifurcation.

As the interpopulation delay *τ* was varied, the two foci traced trajectories in (*G*, *ω*) plane, rotating approximately around a common centre. For small *τ* the in-phase focus had a lower *G*. As *τ* increased, at some point the two foci shared the same *G* and simulations showed suppression of oscillations. Beyond this point, the antiphase focus took over as the one with the lower *G*, reversing the character of the emerging oscillations. This behaviour repeated approximately periodically with increasing *τ*.

### Phenomenological model of phase difference dynamics

To investigate the characteristic arc-shaped distributions observed in the ISPC-PRI plane, we modelled the phase difference *θ*(*t*) as a mean-reverting stochastic process. Phase difference is a circular variable, but since strong attraction to 0 or π keeps trajectories near these values, we approximated the dynamics with a linear Ornstein-Uhlenbeck process:

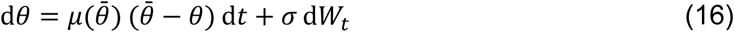

where *θ*^ˉ^ was a target phase difference drawn independently from a uniform distribution over [0, *π*] and fixed for each simulation, *μ* was a target-dependent mean-reversion speed, *σ* was the noise amplitude, and d*W_t_* was the standard Wiener increment. The key assumption encoded in *μ*(*θ*^ˉ^ ) was that mean reversion is fastest when the target phase difference is near 0 or π. Thus, the system was more strongly attracted to in-phase and antiphase relationships than to intermediate phase differences. We used, *μ*(*θ*^ˉ^ ) = *μ*_0_[1 − (2*θ*^ˉ^ /π − 1)^4^]^−1/2^, as an implementation of this assumption; the qualitative results did not depend on the details of this functional form. Euler-Maruyama integration was used (Δ*t* = 0.0001 s) over a window of 32 s. In the first set of simulations, base values of *μ*₀ = 5 s⁻¹ and *σ* = 4 rad·s⁻^1/2^ were used, and 1000 independent realisations were generated. For the second set of simulations, the mean-reversion speed was increased to *μ*₀ = 40 s⁻¹, with noise level adjusted to *σ* = 11.3 rad·s⁻^1/2^ to keep *σ*²/*μ*₀ (and therefore the stationary variance of the process) constant. The two sets of simulations therefore differed in how fast electrode pairs approach the preferred phase difference, while keeping the same long-run phase variability. **Fig. S8** shows the ISPC and PRI values based on these simulations.

## Acknowledgements

We thank Zohreh Zakeri, Neil Mansfield and Caroline Sunderland for contributions to experimental design, data collection, and data preprocessing. AO is grateful to Karen Blackmon for early discussions about the distance dependence of brain synchrony.

## Supplementary Materials

**Fig. S1.**
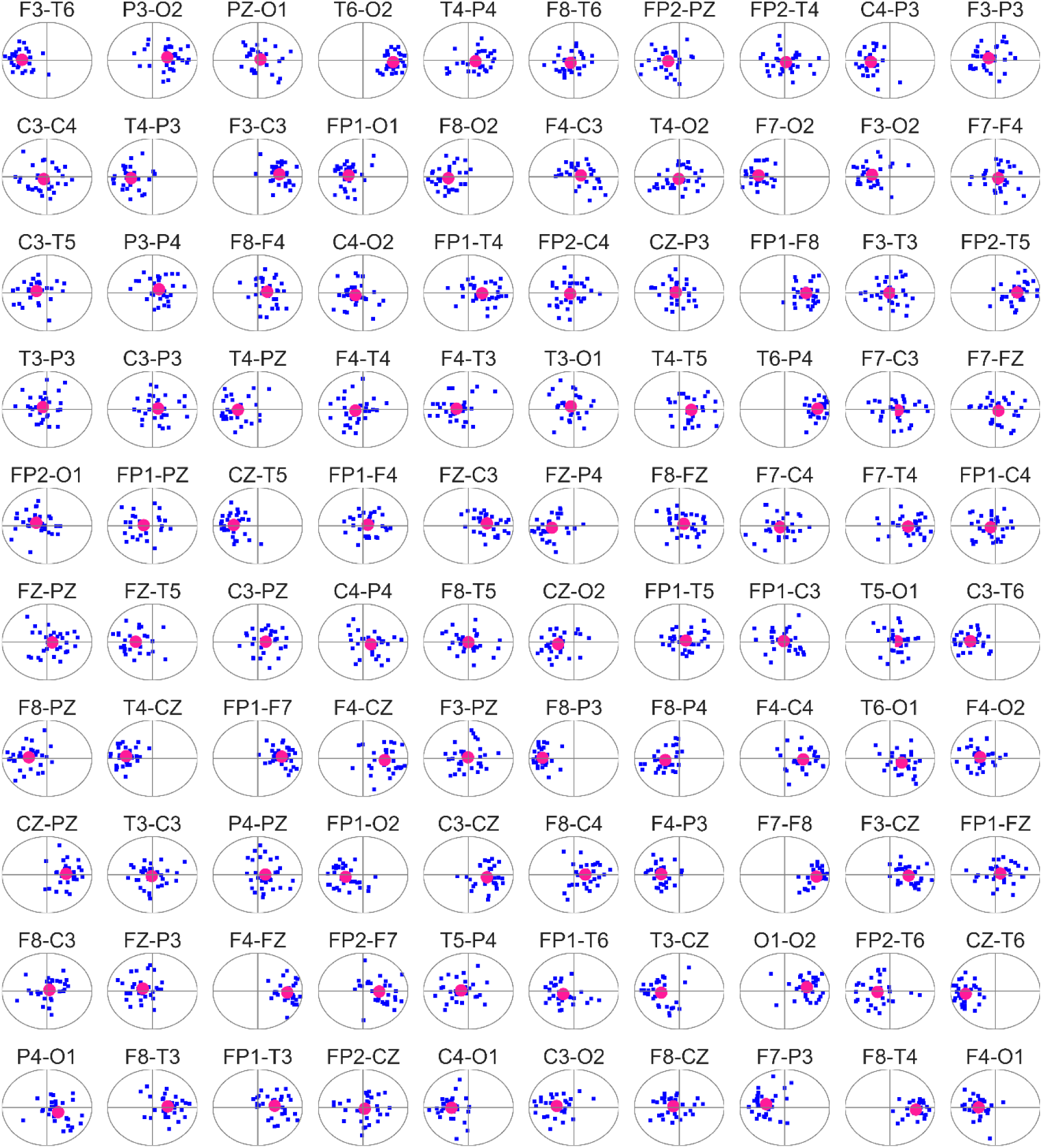
Examples of phase clustering. Type and extent of phase clustering in randomly selected representative electrode pairs (labelled at the top of each subplot) from one participant during a 2 min resting-state EEG, narrow-band filtered at 16 Hz (1 Hz bandwidth). For each pair, the inter-site phase difference vector *z* was computed in adjacent windows of size *W* = 4 s, yielding 30 values (blue dots) with their mean (red circle) shown in the Argand plane with the unit circle indicated in grey.

**Fig. S2.**
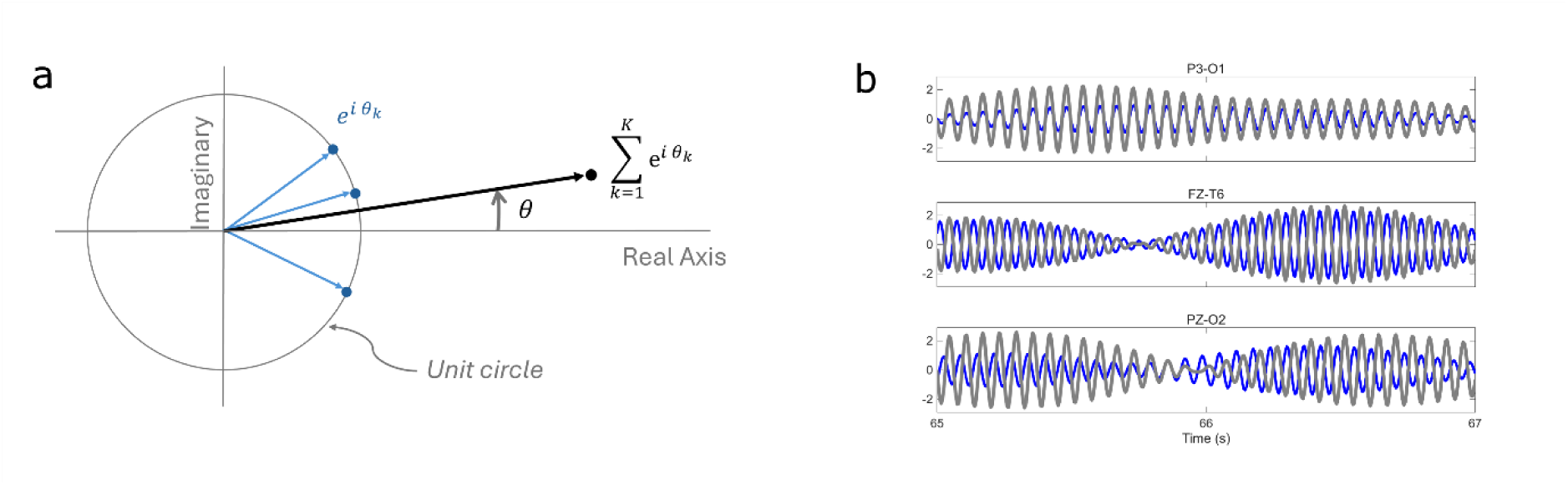
Schematic of phase vector in Argand plane and examples of pairs of EEG signals. **a**, Phase vectors for data samples (blue dots on the unit circle) and their sum (black dot). **b**, A two-second segment of EEG signals (blue and grey) from selected electrode pairs (labelled at top of each subplot) from a representative participant narrow-band filtered at 16 Hz (1 Hz bandwidth), exemplifying characteristic in-phase (top), antiphase (middle) and weak (bottom) clustering.

**Fig. S3.**
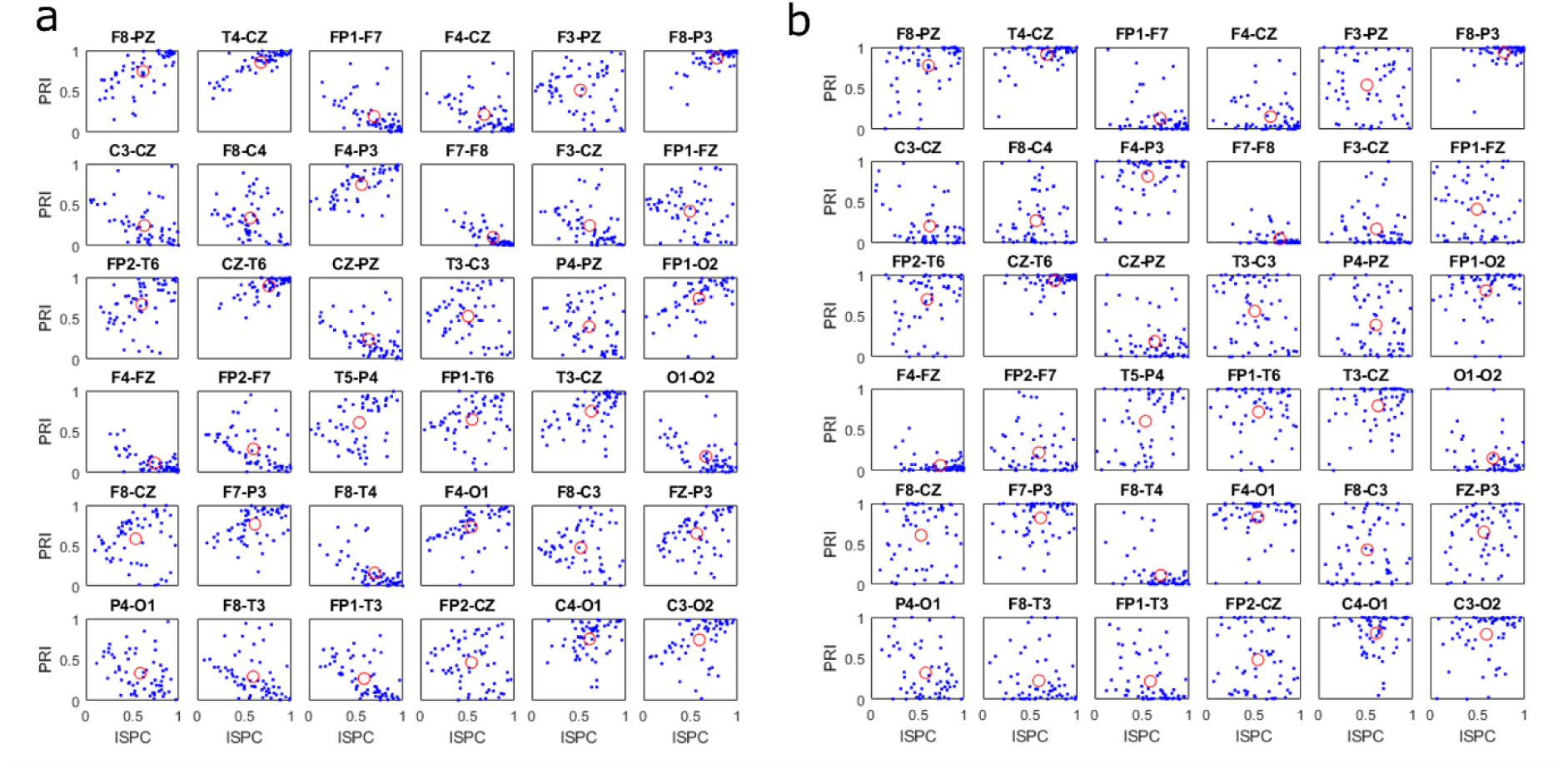
Comparing the results of Hilbert transform and zero-crossing methods for extracting phase. **a**, ISPC and PRI for a representative subset of electrode pairs calculated using the zero-crossing method to extract phases during a 2 min resting-state EEG, narrow-band filtered at 16 Hz (1 Hz bandwidth) in adjacent windows of size *W* = 4 s. **b**, Hilbert transform was used to extract phases.

**Fig. S4.**
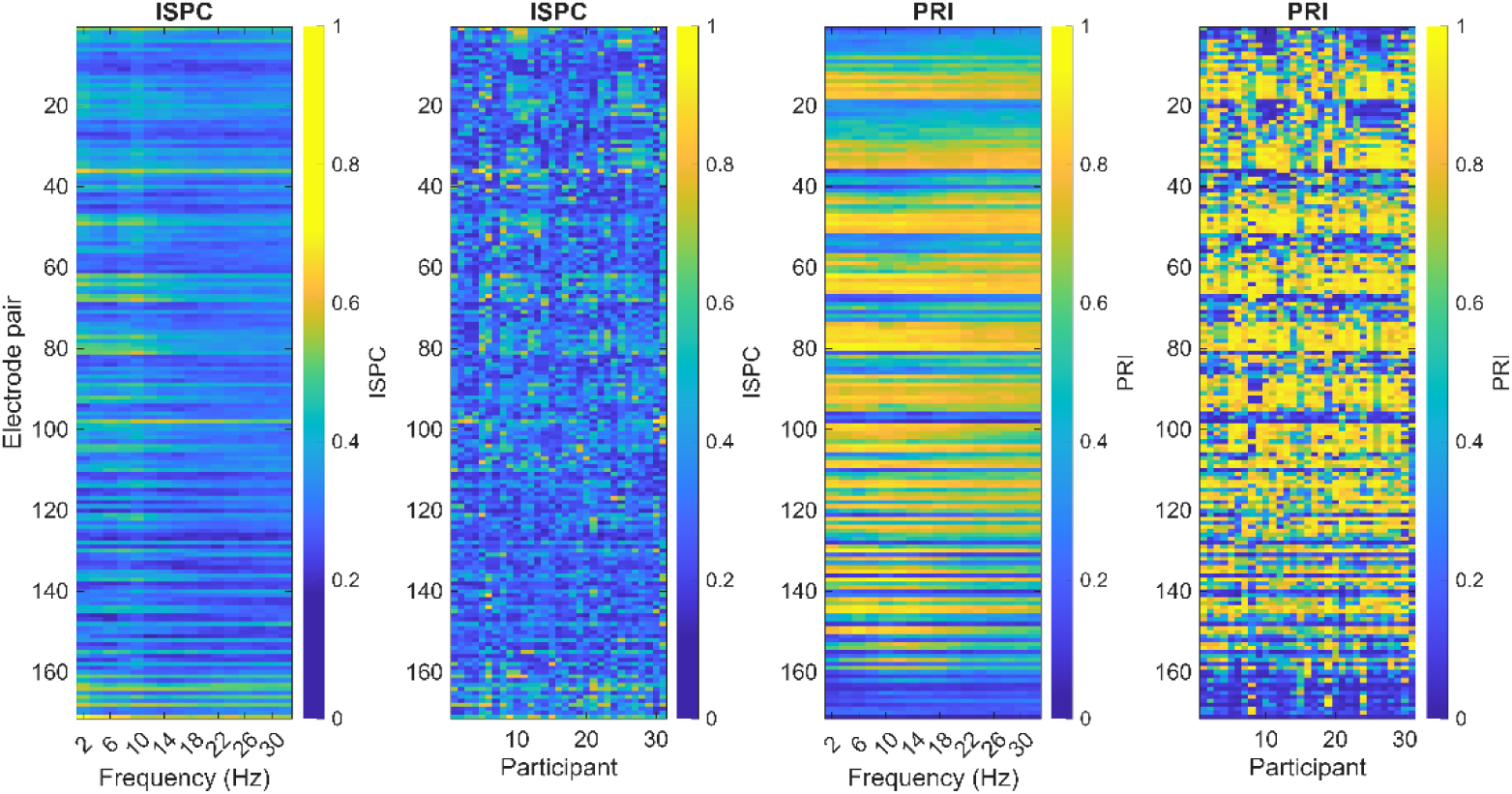
Variability of ISPC and PRI across frequencies and participants. Heatmaps show ISPC (left two panels) and PRI (right two panels) for all 171 electrode pairs (rows) during the initial rest episode (R1). The first and third panels display each metric as a function of frequency (averaged across participants); the second and fourth panels display each metric as a function of participant (averaged across frequencies). Horizontal banding in the frequency panels indicates that both metrics are stable across frequencies for each electrode pair. Greater variation across columns in the participant panels reflects the larger participant-associated variability.

**Fig. S5.**
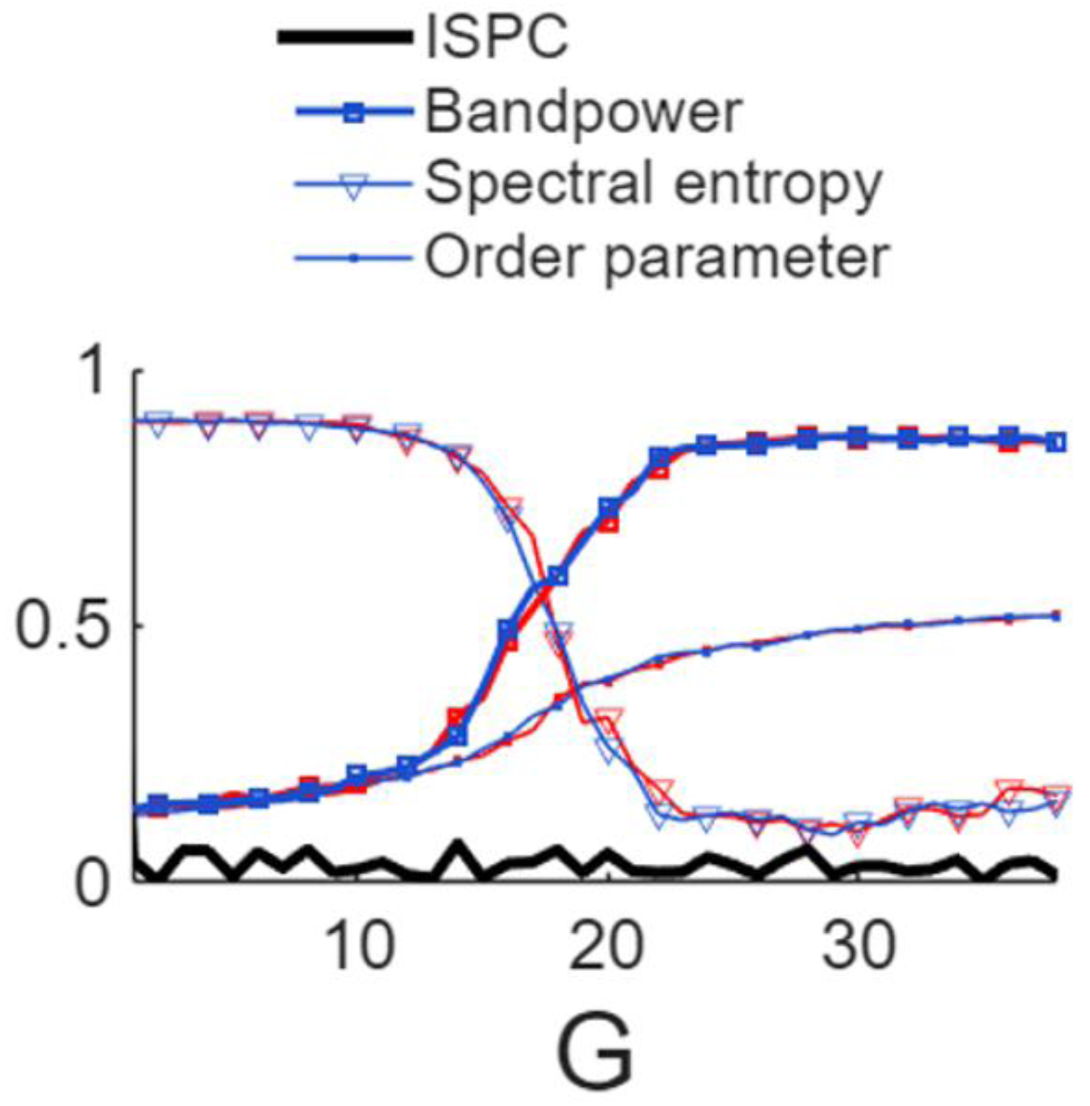
Synchronisation with zero intrapopulation transmission delay (*d* = 0) **and no interpopulation coupling** (*g* = 0) ISPC (black) and three measures of within-population synchronisation, relative band power near the peak frequency (squares), spectral entropy (triangles) and order parameter (dots), are shown for each population (blue, red) as a function of intrapopulation connectivity strength *G*. Entropy values rescaled and shifted for visibility. External input: 1,200 spikes/s per neuron for both populations.

**Fig. S6.**
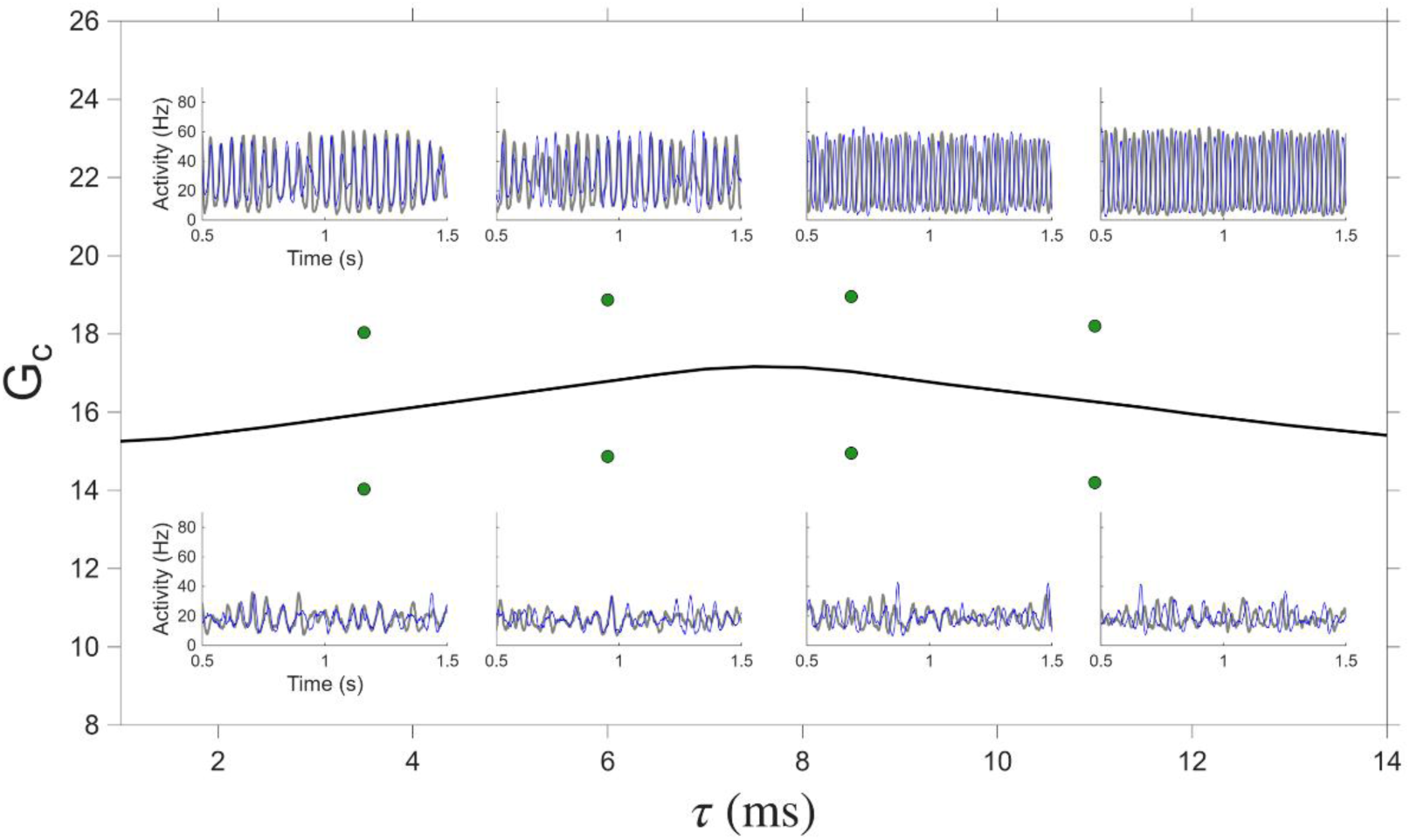
LIF simulation verification of the linear stability analysis. The critical coupling curve *G_c_*(*τ*) (solid black line), reproduced from Fig. 4g, is shown alongside eight sets of simulation parameters (*G*, *τ*) (green dots), four below and four above the curve. Insets show the corresponding population activity timeseries for each parameter set, with blue and grey traces representing the two populations. Dots below *G_c_* yield unsynchronised activity (lower insets), while dots above *G_c_* yield synchronised, phase-clustered activity (upper insets), confirming the predictions of the linear stability analysis. The phase relationship between the two population traces in the upper insets shifts from in-phase (left insets, *τ* < *τ_c_* to antiphase (right insets, *τ* ≥ *τ_c_*), as can be verified by inspecting the relative phase of the blue and grey traces. As *G* approaches *G_c_* from below, transient episodes of synchronised phase-clustered activity emerge with increasing duration until they fully dominate once *G* > *G_c_*. Similarly, as *τ* approaches *τ_c_* from left, transient antiphase episodes appear with increasing duration, eventually replacing in-phase clustering entirely for *τ* > *τ_c_* . Interpopulation gain *g* = 2 ; intrapopulation transmission delay *d* = 1 ms.

**Fig. S7.**
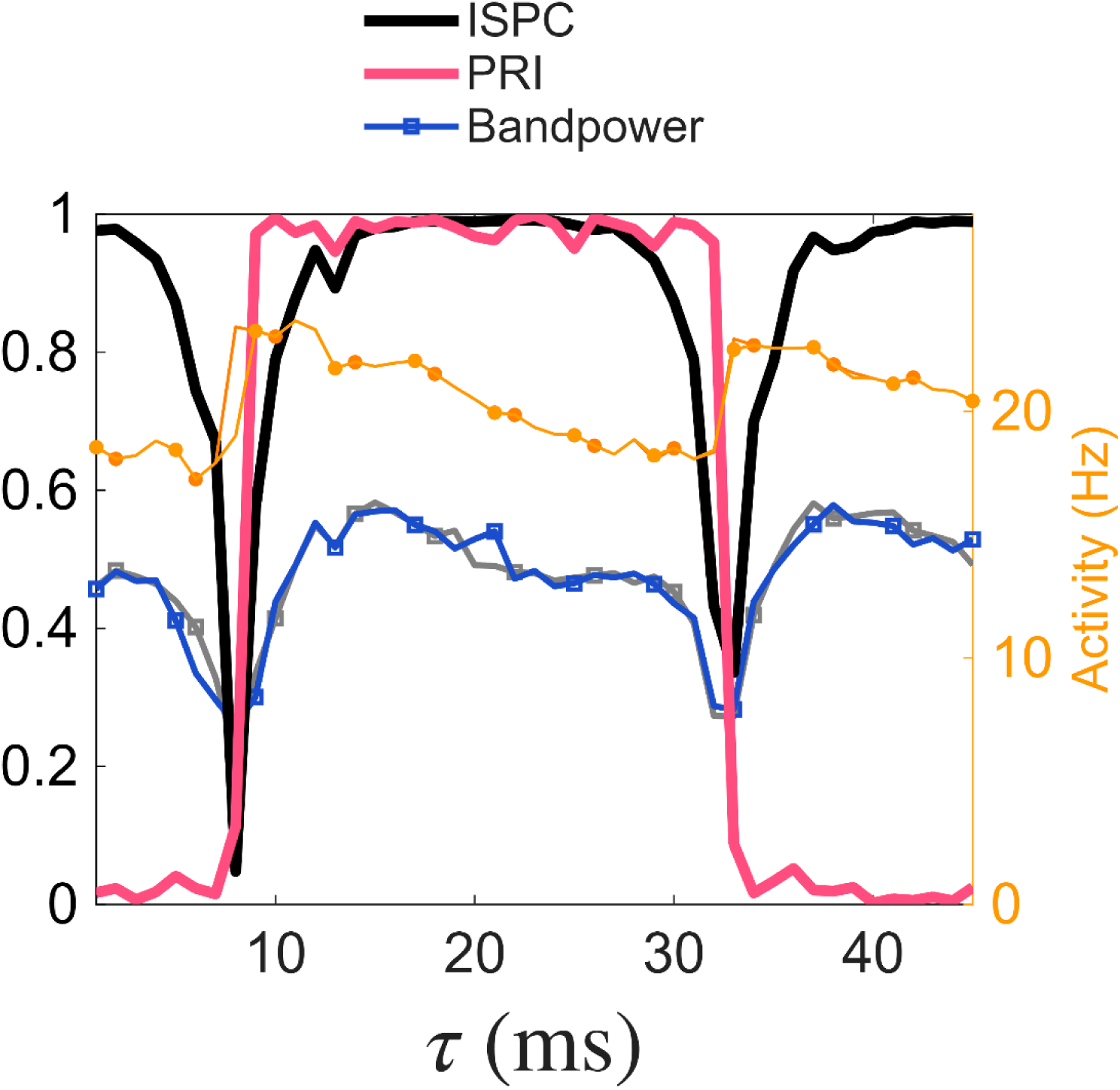
Transitions between in-phase and antiphase modes are approximately periodic in interpopulation delay. As in Fig. 3i but shown over a greater range of interpopulation delay *τ*, illustrating the approximately periodic recurrence of in-phase and antiphase regimes.

**Fig. S8.**
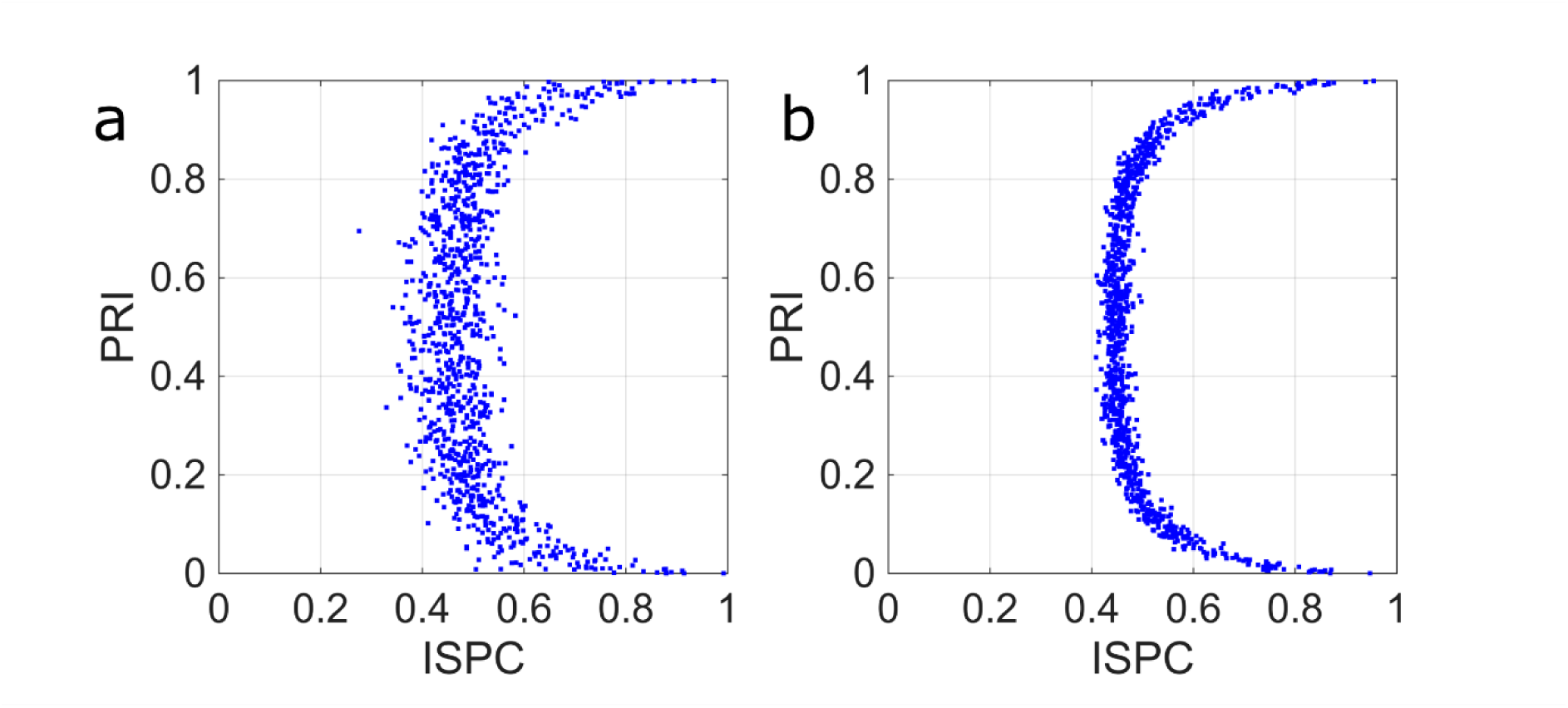
Phenomenological stochastic phase model replicates the arc-shaped ISPC-PRI distribution and its task-evoked tightening. Each panel shows 1000 simulated realisations plotted in the ISPC-PRI plane, where each realisation used an independently drawn target phase difference *θ*^ˉ^ ∼ Uniform[0, π]. In both panels the arc-shaped distribution characteristic of the EEG data is reproduced. **a**, Base parameter regime: *μ*₀ = 5 s⁻¹, *σ* = 4 rad·s⁻^1/2^. **b**, Increased mean-reversion speed: *μ*₀ = 40 s⁻¹, with *σ*²/*μ*₀ held constant across panels so that the stationary variance of the process is the same in both conditions. The tighter arc in (b) illustrates how increasing the mean-reversion speed, without changing the stationary phase variance or the preferred phase differences, reproduces the task-evoked tightening of the arc observed in the EEG data (Fig. 5g-h). See Methods (Phenomenological model of phase difference dynamics) for full model specification.

**Fig. S9.**
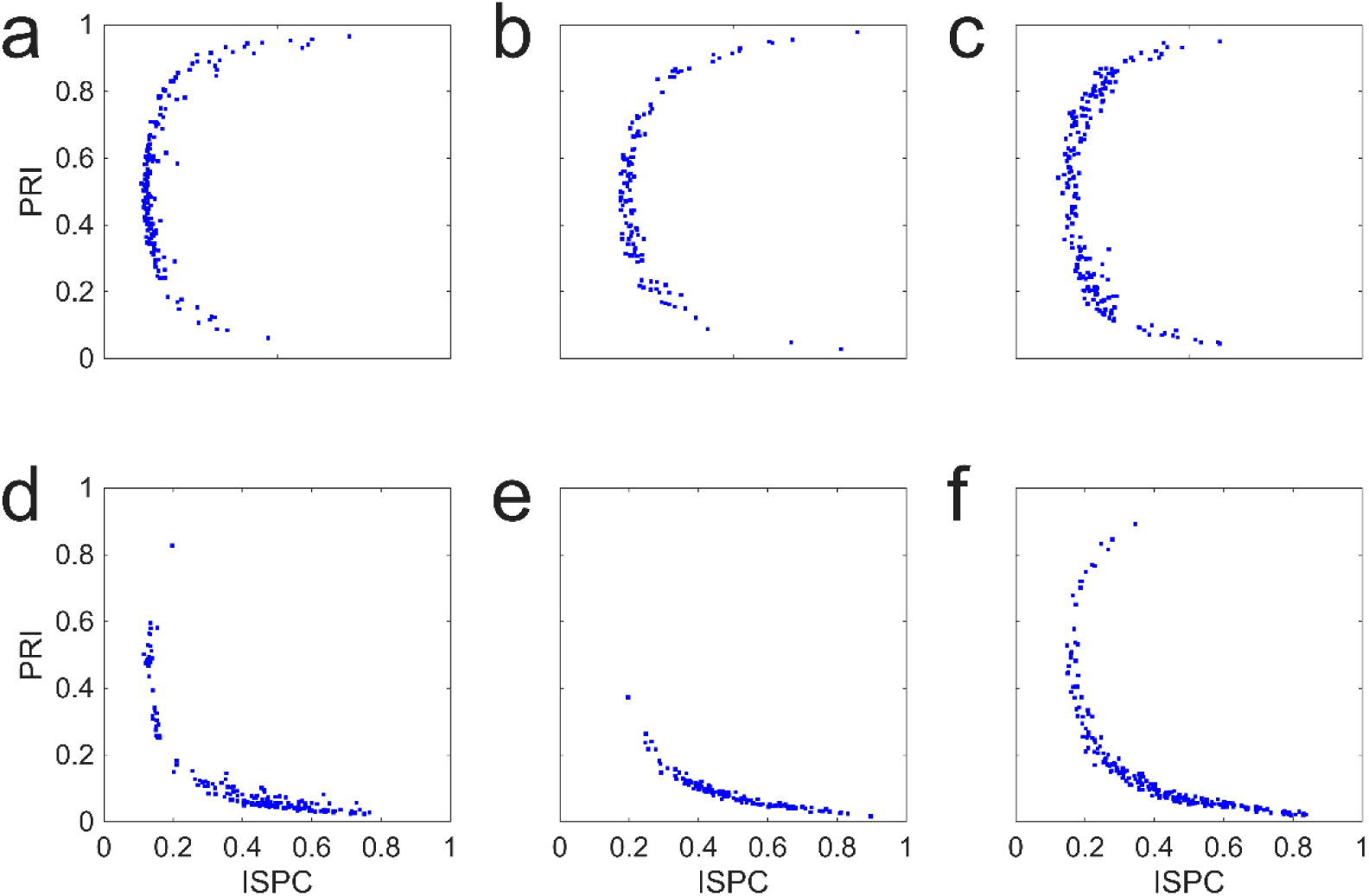
Generalisability of ISPC-PRI distributions across independent datasets and effect of Laplacian spatial filtering. Joint distribution of ISPC and PRI for one representative participant from each of three independent EEG datasets, with (a-c) and without (d-f) surface Laplacian spatial filtering. ISPC and PRI were calculated for each electrode pair (blue dots) after narrow band (1 Hz wide) filtering centred at 16 Hz. **a, d,** 25-month-old healthy child (*W* = 6 s)^36^. **b, e,** Healthy athlete (*W* = 26 s)^35^. **c, f,** 87-year-old participant (*W* = 26 s)^34^. All EEGs had been interpreted by clinical neurophysiologists as normal. Arc-shaped ISPC-PRI distribution with bimodal PRI characteristic of the main dataset (Fig. 1d) are evident across all three datasets when Laplacian filtering is applied (top row). Without Laplacian filtering (bottom row), electrode pairs cluster near PRI = 0 with high ISPC, consistent with the influence of volume-conduction artefact.

**Fig. S10.**
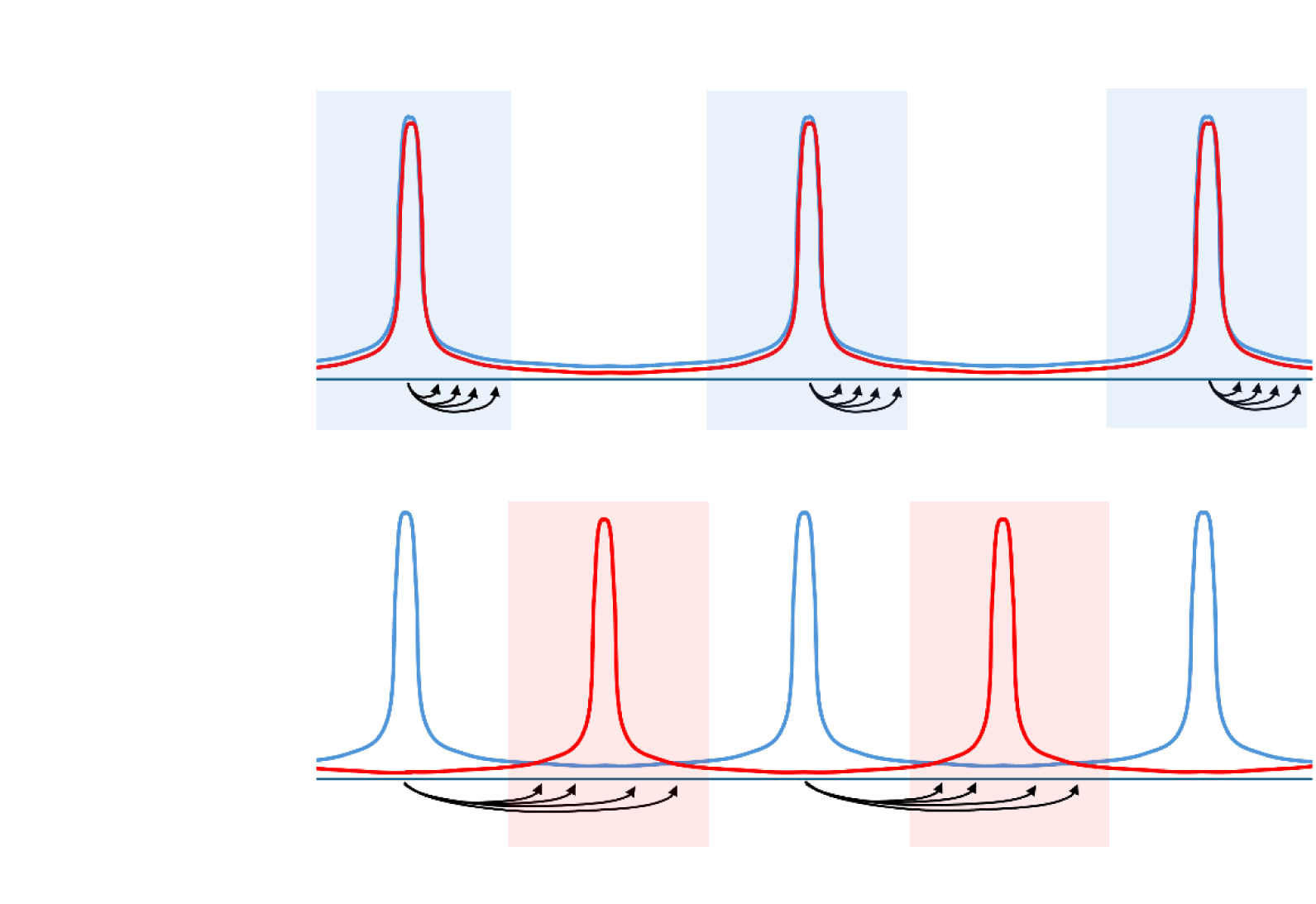
Mutual reinforcement of population spikes in in-phase and antiphase regimes. Schematic illustration of timing relationships for the population spikes of two delay-coupled populations (blue and red). Top row: in-phase oscillation. When the interpopulation delay *τ* is short (less than one quarter of the period *P*), spikes from each population arrive while the other is in its excitable phase (shaded regions), reinforcing synchronous firing. Bottom row: antiphase oscillation. When the delay is between one quarter and three quarters of the period, spikes arrive during the other population’s excitable phase, reinforcing the half-period offset. Simulations indicate that when the interpopulation delay is within these ranges, i.e. *τ* < *P*/4 or *P*/4 < *τ* < 3*P*/4, the populations settle into precise in-phase or antiphase relationships, respectively (Fig. 3d,e,i). Arrows in the figure symbolise excitatory input from the blue to the red population; input from red to blue is omitted for visual clarity.

**Table S1.** Distribution of electrode pairs by cortical distance and connection type. Electrode pairs were grouped into four distance bins (in mm) and categorised as homologous interhemispheric (symmetric pairs across hemispheres), non-homologous interhemispheric (asymmetric pairs across hemispheres), or intrahemispheric (within the same hemisphere). N pairs indicates the number of electrode pairs in each category. Mean, Min, and Max indicate the cortical distance (in mm) within each group. Within each distance bin, the three categories have comparable mean distances, allowing comparison of phase relationships across connection types at matched distances.

| Bin | Distance (mm) | Category | N pairs | Mean | Min | Max |
| --- | --- | --- | --- | --- | --- | --- |
| 1 | 44.7–77.2 | Homologous | 3 | 58.8 | 49.7 | 76.8 |
| 1 | 44.7–77.2 | Non-Homologous Interhemispheric | 0 | N/A | N/A | N/A |
| 1 | 44.7–77.2 | Intrahemispheric | 28 | 57.2 | 44.7 | 76.7 |
| 2 | 77.2–109.6 | Homologous | 2 | 91.9 | 82.1 | 101.7 |
| 2 | 77.2–109.6 | Non-Homologous Interhemispheric | 14 | 94.3 | 79.2 | 107.3 |
| 2 | 77.2–109.6 | Intrahemispheric | 14 | 96.7 | 78.4 | 106.6 |
| 3 | 109.6–142.1 | Homologous | 3 | 124.6 | 116.1 | 140.8 |
| 3 | 109.6–142.1 | Non-Homologous Interhemispheric | 26 | 129.4 | 109.8 | 138.7 |
| 3 | 109.6–142.1 | Intrahemispheric | 10 | 128.1 | 109.9 | 141.6 |
| 4 | 142.1–174.5 | Homologous | 0 | N/A | N/A | N/A |
| 4 | 142.1–174.5 | Non-Homologous Interhemispheric | 16 | 155.8 | 147.2 | 174.5 |
| 4 | 142.1–174.5 | Intrahemispheric | 4 | 154.8 | 142.3 | 166.7 |

**Video S1. Stability analysis of two coupled populations and ongoing simulation showing the switch from in-phase to antiphase behaviour with increasing interpopulation delay**

Results of the repeated linear stability analysis (top row) with continually updated interpopulation delay *τ* are shown together with ongoing simulation results (bottom row). Top left panel displays the unstable foci in the *G*-*ω* plane with the dashed line indicating *G_crit_* (see Methods). In-phase and antiphase foci are indicated by white and red dots, respectively. With increasing *τ*, the foci rotate clockwise around an approximate common centre where the focus with the lower *G* determins the type of instability. Top right panel shows, for the current value of *τ*, the perturbations away from equilibrium for the two populations, *ρ*_1_ (blue) and *ρ*′_1_ (grey), extracted from the unstable eigenfunction of the complex stability matrix. Bottom left panel shows the firing rates of the two populations while the bottom right panel displays *ρ*(*v*, *t*) (blue) and *ρ*′(*v*, *t*) (grey). As interpopulation delay increases and the antiphase focus attains dominance (i.e. has lower *G*), the simulation indicates that the oscillations in activity are initially suppressed and, after a transient, switch to antiphase behaviour. The intrapopulation connectivity ( *G* = 18 ) is fixed slightly above the critical value in the simulation, and the interpopulation connectivity *g* = 2, sufficient to engender phase clustering (in- or antiphase depending on *τ*).

